# Bdelloid rotifers use hundreds of horizontally acquired genes against fungal pathogens

**DOI:** 10.1101/2021.09.04.458992

**Authors:** Reuben W. Nowell, Timothy G. Barraclough, Christopher G. Wilson

**Affiliations:** Department of Zoology, University of Oxford; 11a Mansfield Road, Oxford, OX1 3SZ, UK; Department of Life Sciences, Imperial College London; Silwood Park Campus, Buckhurst Road, Ascot, Berkshire SL5 7PY, UK

## Abstract

Obligately asexual lineages are typically rare and short-lived. According to one hypothesis, they adapt too slowly to withstand relentlessly coevolving pathogens. Bdelloid rotifers seem to have avoided this fate, by enduring millions of years without males or sex. We investigated whether bdelloids’ unusual capacity to acquire non-metazoan genes horizontally has enhanced their resistance to pathogens. We found that horizontally transferred genes are three times more likely than native genes to be upregulated in response to a natural fungal pathogen. This enrichment was twofold stronger than that elicited by a physical stressor (desiccation), and the genes showed little overlap. Among hundreds of upregulated non-metazoan genes were RNA ligases putatively involved in resisting fungal toxins and glucanases predicted to bind to fungal cell walls, acquired from bacteria. Our results provide evidence that bdelloids mitigate a predicted challenge of long-term asexuality in part through their ability to acquire and deploy so many foreign genes.

## Introduction

The maintenance of sexual reproduction is a longstanding problem in evolutionary biology^1–3^. In theory, sexual populations ought to be replaced by asexual competitors, which can reproduce up to twice as efficiently by investing resources in independently reproductive females^4,5^, versus males that make no contribution to growth. Despite this and other evolutionary advantages of asexuality^6^, biparental meiotic sex is nearly universal among eukaryotes^7,8^. Less than 1% of animal and plant species are obligately asexual, and almost all are of recent origin, implying that sex brings such major benefits that clonal lineages are driven extinct^9,10^.

Many genetic and ecological hypotheses have been proposed to explain the prevalence of sex^11,12^, but none has decisive support. One prominent idea is that shuffling genes through sex facilitates adaptation to the intense, ubiquitous and relentlessly changing selection imposed by coevolving parasites and pathogens^13–15^. This ‘Red Queen Hypothesis’ (RQH) has broad theoretical appeal as a mechanism to favour genetic mixing and suppress asexuality, either by itself^16,17^ or in combination with other processes^18–20^, especially when pathogens are highly virulent^21^. It is hard, however, to find appropriate systems to test this hypothesis in nature^22,23^. Cases where sexuals and obligate asexuals coexist offer some support, but these are unusual^6^, and even studies lasting many years do not necessarily reflect stable evolutionary outcomes^24^. Given that asexuals can outcompete sexual relatives locally or temporarily^25^, and many predicted fitness costs of clonality are not easily measured^26^, a longer view is vital for understanding why they almost inevitably disappear.

One approach is to investigate taxa where males and sex seem to have been absent for extended evolutionary timescales^27^. Whatever mechanisms maintain sex in the majority of organisms ought to be absent, weakened or unusually mitigated in these groups. Understanding these cases can thus shed light on the rules that govern sex and keep clonality in check everywhere else^28^. The RQH posits a key role for coevolving parasites and pathogens. If so, ‘ancient asexuals’ ought to have unusual genetic or ecological mechanisms for dealing with disease, which would be of interest not only in understanding sex, but in helping to test broader models of coevolution, genetic diversity and immunity^29–32^.

Bdelloid rotifers present a useful system for addressing such questions. This class of tiny freshwater invertebrates has persisted for tens of millions of years and diversified into over 450 species^33,34^. Reproduction is only known by parthenogenetic eggs and neither males nor sperm have been reported in contemporary populations, unlike other putatively ancient asexual animals^35–40^. Genetic evidence to either confirm or refute their long-term asexuality has proved complicated to obtain, and evidence previously reported for and against this hypothesis^41–44^ has been overturned by later studies^45–49^. Within individuals, genes occur in multiple copies that appear to share sequence homology via ongoing gene conversion^43,45^ and recombination^50^, but some authors have also proposed unusual modes of genetic exchange between individuals, based on exotic mechanisms unknown in other animals^44,51,52^. While it is difficult to exclude that bdelloids engage in some form of inter-individual recombination that remains obscure^50^, they nevertheless comprise an exceptional animal clade in which conventional sex is so rare or cryptic as to elude centuries of observation^53,54^ and remain opaque to direct study.

If the RQH helps explain the prevalence of sex, we might expect that bdelloids either lack virulent pathogens, or possess unusual characteristics that mitigate the impact of these natural enemies. In fact, bdelloids are attacked by over 60 species of virulent fungal (e.g. **Fig. 1a**) and oomycete pathogens that specialize on them^55^. These can exterminate cultured populations in a few weeks^56,57^, and significantly depress the abundance of hosts in natural habitats^58^. Controlled infection trials show substantial variation in resistance and infectivity phenotypes among host and fungal pathogen isolates (**Fig. 1b**; unpublished data), suggesting the possibility of genetically specific mechanisms of immunity whose efficacy varies depending on the host-pathogen pairing, as envisaged by Red Queen models^16,59^ and observed in other invertebrate systems^60–65^. Nothing is currently known, however, about the genetic basis of these resistance traits in bdelloids (or rotifers more generally), or how they might evolve and remain effective in the absence of sex.

**Fig. 1.**
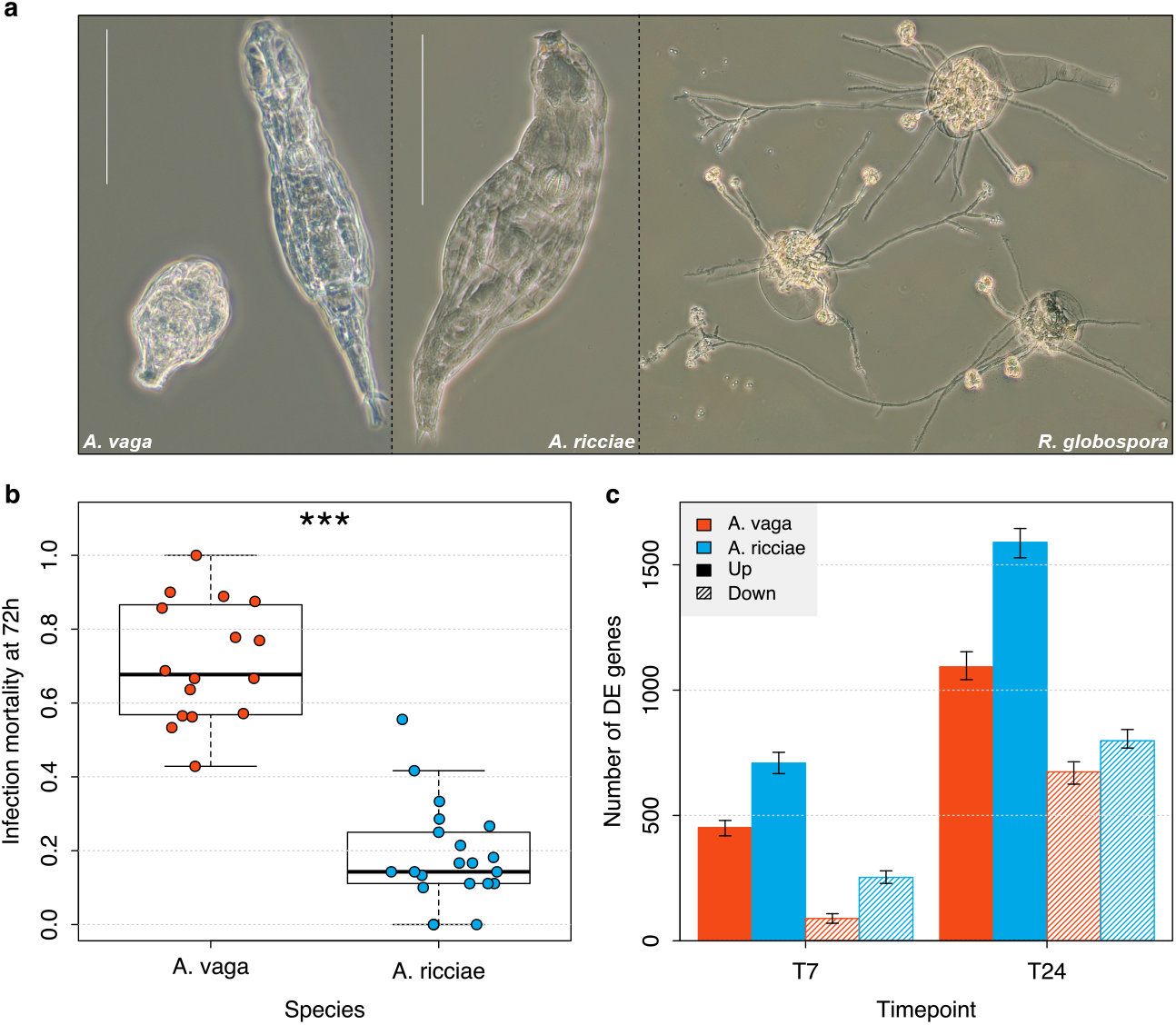
The response of bdelloid rotifers to experimental inoculation with *R. globospora*. **a.** Active and contracted animals of *A. vaga* (left panel); healthy *A. ricciae* (middle panel); composite image of three *A. ricciae* corpses with fully developed *R. globospora* infections, with hyphae differentiating into irregular resting spores and long conidiophores bearing spherical infectious conidia (scale bars 100μm). **b.** Proportion of rotifers killed by infection 72 hours after initial exposure to *R. globospora*. Points indicate replicate laboratory populations with approximately 10 individuals (range 6 to 20) exposed to 1000 conidia. **c.** Numbers of genes showing significant upregulation (solid bars) and downregulation (hatched bars) at 7- and 24-hours post-inoculation (T7 and T24 respectively), for *A. vaga* (red) and *A. ricciae* (blue), relative to control populations inoculated with UV-inactivated spores.

One potential genetic mechanism that could help bdelloid rotifers diversify and sustain resistance against pathogens is the incorporation of new genes by horizontal gene transfer (HGT). Approximately 10% of genes in bdelloid genomes appear to have been acquired from bacteria, fungi, plants and other non-metazoan sources, rather than sharing recent common ancestry with orthologs in related animal groups^43,51,66^. This estimate is an order of magnitude greater than in other animals^67^, holds for all bdelloid genomes so far examined, and is robust across various approaches to identifying HGT events^46,68–70^.

Comparisons among bdelloid species indicate that most HGT events are ancient but with ongoing rates of stable gene acquisition estimated to be on the order of one gain per 100,000 years^71^. At these rates, the phenomenon is too slow to be equated with the shuffling of alleles in a population by sex, but HGT has been hypothesized to introduce novel biochemical functions that help bdelloids adapt to environmental challenges. Foreign genes are indeed expressed and incorporated into metabolic pathways^66,68^, some of which are not shared by other metazoans^72^. Furthermore, these genes have been found to play a role in desiccation tolerance, nutrient exploitation and repair of DNA damage^66,68,73^. However, the intense and continual selection resulting from evolving pathogens might particularly favour the acquisition and co-option of genes involved in immune responses, such as defensive proteins or antimicrobial compounds. HGT in bacteria is commonly associated with functions and products that help combat antagonists^74,75^ and isolated examples of HGT contributing to immunity are known from invertebrates^76–78^. The massive scale of HGT in bdelloids might therefore have brought new gene functions that could compensate in part for the theoretical challenge that pathogens pose to asexual groups, as posited by the RQH.

To test this prediction, we used RNA-seq to identify genes that are differentially expressed when bdelloid rotifers are attacked by a fungal pathogen that in the genus *Rotiferophthora* (Clavicipitaceae, Hypocreales), which preys specifically upon them^79^. We assessed variation in this response by comparing two species, *Adineta ricciae* and *A. vaga*, which differ in resistance by a factor of four to the same pathogen. We used functional annotation to identify putative defensive or immune-related genes and tested whether these showed evidence of horizontally acquisition from non-metazoan taxa. Finally, we compared our results with RNA-seq data obtained when bdelloids were exposed to desiccation^73^, enabling us to test whether genes of foreign origin contribute disproportionately when responding to a biotic as opposed to an abiotic stressor. We found strong support for the prediction that HGT plays a disproportionate role in defence against pathogens. Hundreds of foreign genes were differentially regulated in response to fungal attack, some producing proteins not previously implicated in immune defence in animals. Our results demonstrate the scale and range of functions that HGT adds to the genetic repertoire of bdelloids, particularly in the important context of adaptation to biotic antagonists.

## Results and Discussion

Clonal populations of the bdelloid rotifers *Adineta ricciae* and *A. vaga* were experimentally inoculated under controlled conditions with spores of the virulent fungal pathogen *Rotiferophthora globospora*^79^, as part of a larger study. In both species, 95% of animals ingested spores and contracted (**Fig. 1a**) within minutes of exposure; they were then monitored over 3–4 days (see Methods). The proportion of animals succumbing to fungal infection differed greatly between the species: 18% for *A. ricciae* and 71% for *A. vaga* (**Fig. 1b**, relative risk = 3.74, 95% CI: 2.8-5.0, z = 8.758, P <0.001). Given that these closely related species are ecologically^46,80^ and morphologically similar (**Fig. 1a**) and were reared under standard conditions, the basis of the difference in mortality seemed to be physiological, reflecting a strong defensive response in *A. ricciae*, and a weaker defence in *A. vaga*. To investigate the genetic basis of this response, we repeated the inoculation experiment at scale, and harvested total RNA from exposed and control populations of each species in triplicate at 7h and 24h, during the early stages of fungal attack (Methods). We took advantage of previously published reference genomes for both species^43,46^ to assemble, map and quantify sequenced transcripts, and compared species and timepoints for patterns of differential gene expression between inoculated and control populations (**Fig. 1c**, Methods).

We defined significant differential expression stringently, as a >4-fold difference in expression level and a False Discovery Rate-adjusted *P*-value < 0.001 (**Fig. 1**, Methods). By these criteria, 541 *A. vaga* genes and 962 *A. ricciae* genes were differentially expressed at 7 hours post-inoculation (T7). Most of these were upregulated in the pathogen treatment group relative to controls (452 in *A. vaga*, ~84%; 709 in *A. ricciae*, ~74%). At 24-hours postinoculation (T24), the number of differentially expressed genes rose to 1767 in *A. vaga* (with 1093 upregulated, ~62%), and 2388 in *A. ricciae* (1590 upregulated, ~67%) (**Fig. 1**). The identities of the differentially regulated genes in each species at T7 overlapped substantially with those at T24, and the direction and magnitude of differential expression at both timepoints was significantly correlated (Pearson’s correlation *R* = 0.66–0.84; *P* < 2e–16 in all cases; Supplementary Fig. 1). This indicates that many of the transcriptional changes initiated at T7 were ongoing and consistent at T24, even as new genes joined the response. In both species, the numbers of differentially expressed genes showed a comparable increase between T7 and T24, but the response in *A. ricciae* was initially stronger and remained higher, especially among upregulated genes. The more rapid and extensive transcriptional response in *A. ricciae* might be a broad-scale physiological reflection of its ability to recognise the pathogen sooner and mobilise defences more extensively than *A. vaga*. The two species showed overlapping profiles of differentially expressed genes, as expected if some defensive or recognition pathways were conserved (Supplementary Fig. 2; Supplementary Table 3). For example, of the 1093 genes that were significantly upregulated in *A. vaga* at T24, 552 (50.5%) shared an ortholog that was also significant upregulated in *A. ricciae*, and these orthologs showed a significant correlation in the magnitude of differential expression (Pearson’s correlation *R* = 0.62, *P* < 2e–16).

Based on the predicted proteins from the reference genomes^46,70^, the background frequency of candidate HGT genes (HGT_C_) in both species is ~11%. Among the significantly upregulated genes, however, the frequency of HGT_C_ increased to 20–30% (**Fig. 2**; **Table 1)**, which represents a 2- to 3-fold enrichment (Fisher’s exact test for an association between upregulation and HGT_C_, *P* < 0.001 in all cases). The pattern was repeated at both timepoints; the proportional enrichment was stronger in the early response, 7 hours after inoculation, but the absolute numbers of upregulated HGT_C_ genes were higher at T24, with 345 genes of putatively foreign origin upregulated in *A. ricciae*, and 269 in *A. vaga*. The proportion of HGT_C_ was not significantly different between the two species (**Table 1**). These results are robust to the threshold of fold-change used to define differential expression (1.5-fold, 2-fold, 8-fold, and 16-fold absolute differences yielded the same results as 4-fold, Supplementary Figs. 3, 4). We also observe a significant but lesser enrichment of HGT_C_ among genes that are significantly downregulated during the experiment (Fisher’s exact test, *P* < 0.001 in all cases; **Fig. 2**; **Table 1**; Supplementary Figs. 3, 4). Horizontally acquired genes were therefore significantly overrepresented among the genes that are differentially expressed, and especially upregulated, in the infected treatments. Comparison of control populations (Supplementary Figs. 5, 6) showed that HGT_C_ genes are more likely to be expressed than native genes under control conditions, but at significantly lower levels. The enrichment of HGT_C_ among differentially expressed gene is thus unlikely to be explained by a known false-positive bias in some RNA-seq analyses toward genes with higher expression levels^81,82^.

**Fig. 2.**
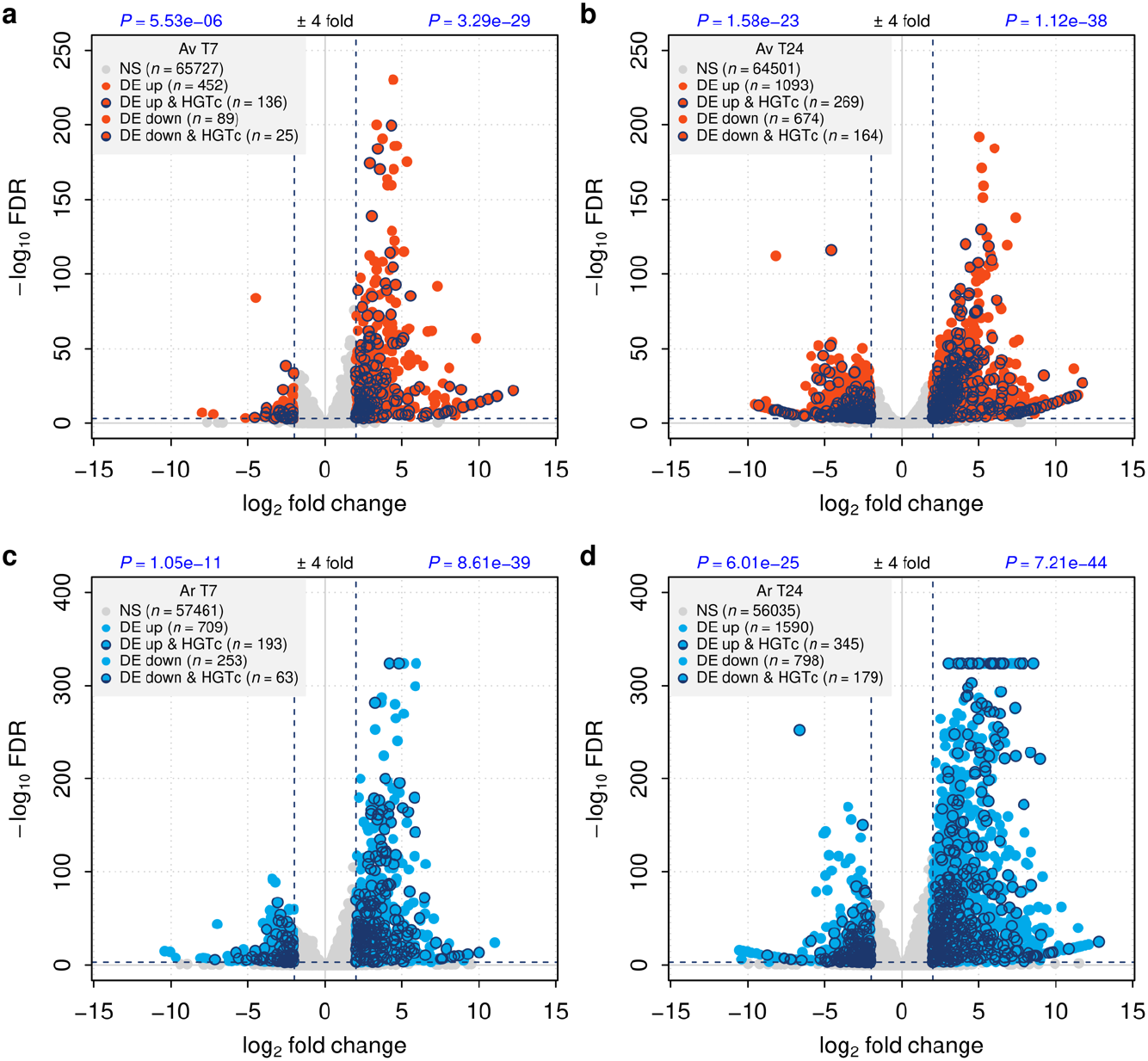
Up- and downregulation of native and foreign genes in response to fungal infection in two species of bdelloid rotifer. Data are shown for **a.** *A. vaga* T7; **b.** *A. vaga* T24; **c.** *A. ricciae* T7 and **d.** *A. ricciae* T24. Each point on the ‘volcano’ plot represents a gene plotted by log_2_ fold-change in expression level on the *X*-axis, and significance level (expressed as –log_10_ FDR) on the *Y*-axis. Positive *X*-axis values indicate genes that were upregulated in the treatment groups (i.e., those exposed to the live pathogen) relative to the control groups, whereas negative values indicate genes that were downregulated in the treatment group relative to the control group. Genes with significant changes in expression level (absolute fold-change > 4 and FDR < 1e–3, dashed lines) are shown in colours (red: *A. vaga;* blue: *A. ricciae);* non-significant changes in expression are indicated with grey (note that the majority of genes are non-significant at these thresholds; see legends for details). *P*-values in blue show the probability of non-association between HGT_C_ and the corresponding up- or downregulated subset (Fisher exact tests, see **Table 1** for details).

**Table 1.** Proportion of foreign genes in significantly up- and down-regulated gene sets during pathogen exposure and entry/recovery from desiccation.

| Test<sup>1</sup> | Species | Contrast | DE type<sup>2</sup> | %HGT in DE0<sup>3</sup> | %HGT in DE1 | Odds ratio (95%-CI) | P-value<sup>4</sup> |
| --- | --- | --- | --- | --- | --- | --- | --- |
| Pathogen | *A. vaga* | T7 | Up | 10.5% | 30.1% | 3.6 (2.9–4.4) | <0.001 |
|  |  |  | Down | 10.6% | 28.1% | 3.2 (1.9–5.2) | <0.001 |
|  |  | T24 | Up | 10.4% | 24.6% | 2.8 (2.4–3.2) | <0.001 |
|  |  |  | Down | 10.5% | 24.3% | 2.7 (2.2–3.2) | <0.001 |
|  | *A. ricciae* | T7 | Up | 9.8% | 27.2% | 3.4 (2.9–4.1) | <0.001 |
|  |  |  | Down | 10.0% | 24.9% | 3.0 (2.2–4.0) | <0.001 |
|  |  | T24 | Up | 9.7% | 21.7% | 2.6 (2.3–2.9) | <0.001 |
|  |  |  | Down | 9.9% | 22.4% | 2.6 (2.2–3.1) | <0.001 |
| Desiccation | *A. vaga* | Entering | Up | 10.6% | 13.1% | 1.2 (1.0–1.6) | 0.06 |
|  |  |  | Down | 10.7% | 9.4% | 0.9 (0.6–1.2) | 0.49 |
|  |  | Recovering | Up | 10.6% | 14.2% | 1.4 (1.2–1.6) | <0.001 |
|  |  |  | Down | 10.5% | 19.1% | 2.0 (1.7–2.3) | <0.001 |

<sup>1</sup>“Pathogen”: analysis of DE where treatment represents animals exposed to live pathogen spores, versus controls exposed to inactivated spores, at 7h (T7) and 24h (T24) post exposure. “Desiccation”: analysis of DE where
treatment represents animals either entering into or recovering from desiccation, versus control animals that remained hydrated (Hecox-Lea & Mark Welch 2018). <sup>2</sup>“Up”: genes that are upregulated in the treatment group; “Down”, genes that are downregulated in the treatment group. <sup>3</sup>“%HGT<sub>C</sub> in DE0”: the proportion of HGT<sub>C</sub> among genes that are not significantly differentially expressed, either up or down depending on DE type; “%HGT<sub>C</sub> in DE1”: the proportion of HGT<sub>C</sub> among genes which are significantly differentially expressed. Significant DE is defined as genes with >4-fold absolute change in expression and FDR < 1e-3. <sup>4</sup>Fisher’s exact test for a significant association between classifications; null hypothesis: true odds ratio not equal to 1.

We performed further analyses to investigate whether enrichment of HGT_C_ genes is specific to the response to fungal infection, or a more widespread phenomenon among differentially expressed bdelloid genes. For example, HGT_C_ genes might be more likely to be differentially expressed in general if they are overrepresented among effectors at the peripheral ends of gene regulatory networks or if their expression is less tightly regulated than that of typical host genes^83,84^, or systematically higher, leading to read-count bias^81^. Finally, HGT_C_ genes might be overrepresented among general stress-response genes, rather than associated specifically with the response to a pathogen; for instance, evidence from earlier analyses has suggested a role for such genes in DNA repair, detoxification and desiccation tolerance in bdelloids^66,68,73^.

We therefore repeated our analyses on published RNA-seq data from a recent study^73^ that investigated gene expression in the same cultured strain of *A. vaga* responding to desiccation. Most bdelloid species are remarkably tolerant to drying out, entering a state of physiological dormancy known as anhydrobiosis^85,86^. ecox-Lea and Mark Welch (2018) compared levels of gene expression in hydrated animals to those that were either entering desiccation or recovering from a period of dormancy induced by desiccation. Thus, we were able to test whether HGT_C_ genes are also enriched in the response to desiccation, using the same criteria applied above for differential expression during infection. Overall, the proportion of HGT_C_ genes involved in the desiccation response was substantially lower than that seen in the pathogen experiment (~9–19%, depending on the subset, compared to ~22–30% previously; **Table 1**). There was a significant enrichment of HGT_C_ when animals were recovering from desiccation (Fisher’s exact test, *P* < 0.001; **Table 1**; Supplementary Fig. 7), but not when animals were entering desiccation. Thus, HGT candidates appear to be disproportionately represented among genes responding strongly to both kinds of stresses, but the effect is ~2-fold stronger during the response to pathogens compared with desiccation, consistent with a more prominent role for HGT_C_ in immune defence.

Next, we compared the identities of genes that were differentially expressed in the fungal infection and desiccation experiments, to test how far these responses are specific to each challenge, rather than reflecting general stress response pathways in *A. vaga* common to both treatments. In general, the proportion of upregulated genes shared between any two sample sets was low (mean = ~26%; Supplementary Table 4), with the exceptions of T7 versus T24 in the infection experiment (~90% of T7 genes also were upregulated at T24), and entering versus recovering from desiccation (~63% of ‘entering’ genes also were upregulated in ‘recovering’). HGT_C_ genes showed a similar pattern. For example, of the 256 HGT candidates upregulated during recovery from desiccation, only 31 (~12%, T7) and 61 (~24%, T24) were also upregulated during pathogen response (Supplementary Table 4). Even fewer genes were shared between downregulated subsets (none and 24 for T7 and T24, respectively). The majority of differentially expressed HGT_C_ in *A. vaga* therefore are unique to either the desiccation or infection experiment.

Having identified hundreds of differentially expressed HGT_C_ genes that appear to be associated specifically with infection, we investigated putative functions by annotating the translated transcriptomes of *A. vaga* and *A. ricciae* based on the presence of Pfam and InterPro domains (see Methods), and found several proteins of particular interest. In *A. vaga*, at T7 and T24, the two foreign genes with the strongest evidence of upregulation are homologous copies of each other, encoding putative RNA ligase (Pfam accession PF13563) and endo/exonuclease domains (PF03372). A further 16 HGT_C_ genes with putative RNA ligase activity (including PF09414 and PF09511) were significantly upregulated across either timepoint. A similar pattern was seen in *A. ricciae*, and gene ontology (GO) analysis confirmed enrichment of RNA ligase/repair activity for both species. Phylogenetic analysis of rotifer-encoded RNA ligase domains supported the classification of these genes as HGT_C_, showing high support for the clustering of rotifer copies with non-metazoan lineages (including bacteria, fungi, viruses, and non-metazoan eukaryotes; **Fig. 3a–c**). The hypothesis that these sequences arise from contamination is refuted by the presence of corresponding paired copies on genomic scaffolds for *A. vaga* and *A. ricciae* whose divergence matches the rest of the genomes, by the fact that these genes are quite distinct from the nearest sequenced non-metazoan copies, and by the presence of introns in genes encoding proteins whose closest homology is otherwise to bacteria. Many upregulated gene copies were closely related to each other and showed expected patterns of homologous, homoeologous or orthologous relationships given the ancestral tetraploidy in bdelloid species^45,87^.

**Fig. 3.**
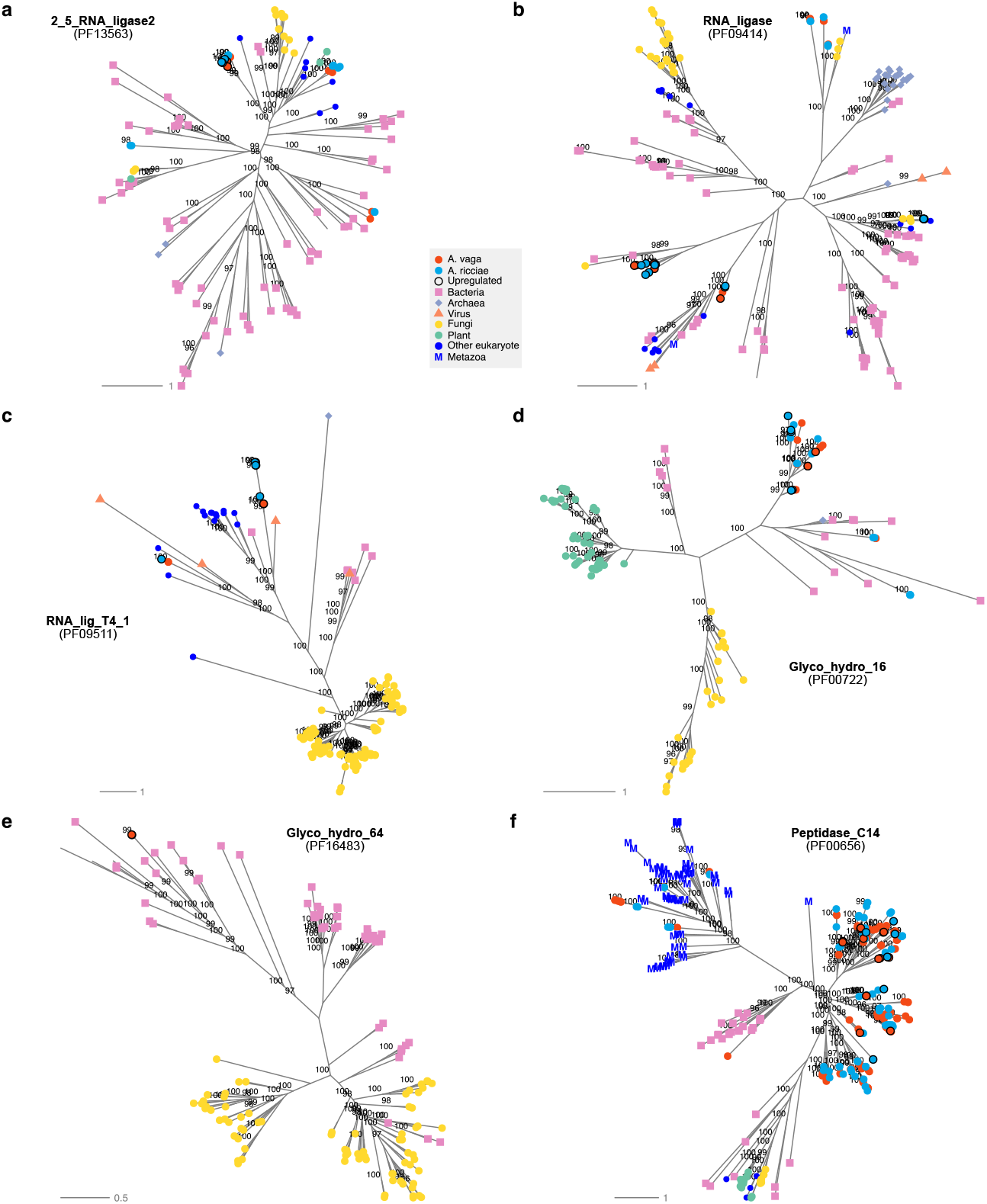
Phylogenetic placement of foreign genes with putative roles in immune function in bdelloids. Phylogenies are shown for the RNA ligase domains **a.** 2_5_RNA_ligase2, **b.** RNA_ligase and **c.** RNA_lig_T4_1; the glycosyl hydrolase domains **d.** Glyco_hydro_16 and **e.** Glyco_hydro_64; and **f.** the caspase domain Peptidase_C14. Pfam accessions are in parentheses. Sequences from *A. vaga* and *A. ricciae* are shown in red and blue, respectively; genes that were upregulated on exposure to the fungal pathogen are highlighted with a black circle. Non-rotifer sequences were taken from the Pfam seed alignments for each domain, and thus represent the known diversity across the tree of life. Tip symbols indicate taxonomic classifications (see legend). Metazoan sequences are indicated with a blue ‘M’ if present.

While the precise function of these genes is unknown, a role for RNA ligase activity in the response of bdelloids to fungal infection has intriguing links to work in other systems. Two close relatives of *Rotiferopthora* in the fungal order Hypocreales attack their insect hosts using lethal ribotoxins as virulence factors^88–91^. These secreted ribonucleases cleave the critical and highly conserved sarcin-ricin loop in the large ribosomal subunit of host cells, which inhibits protein synthesis and causes death^89^. If *Rotiferopthora* hyphae secrete similar virulence factors against *Adineta*, rapid upregulation of proteins with RNA ligase activity might help the host counter this element of the attack by protecting and repairing ribosomes^92^. According to a recent review, RNA toxins, RNA ligases and related RNA repair systems are “extensively disseminated by lateral transfer between distant prokaryotic and microbial eukaryotic lineages consistent with intense inter-organismal conflict”^92^. Might bdelloid rotifers have borrowed machinery from this microbial RNA warfare to defend against their own pathogens? Future work will be needed to explore this speculative hypothesis, by testing whether *R. globospora* expresses such a toxin and determining its effects on rotifer hosts.

Another potential source of immunity to fungi is production of carbohydrate-active enzymes (CAZymes) to target components of fungal cell walls such as chitin and glucans, either acting as recognition receptors or degrading them directly^93–95^. We surveyed 35 glycosyl hydrolase and carbohydrate binding gene families for significant up- or down-regulation, but found no large-scale upregulation of putative CAZymes in response to fungal infection in bdelloids, including genes with putative chitinase and chitin-binding properties. However, a small number of HGT_C_ genes with putative glucanase activity were upregulated in one or both species. These included six genes in both species containing GH16 domains (active on *β*-1,4 or *β*-1,3 glycosidic bonds in various glucans and galactans) with closest matches to the bacterial phylum Bacteroidetes, and two genes restricted to *A. vaga* with glycosyl hydrolase family 64 (GH64, a *β*-1,3-glucanase) domains that cluster with homologs from firmicute bacteria (**Fig. 3d, e**). Glucanases are secreted by plants and bacteria as antifungal compounds^96,97^, and related domains are implicated in insect antifungal defences against *Metarhizium*, a close relative of *Rotiferophthora*^94^. Future work could explore whether these acquired genes play a similar role in bdelloids.

We searched the upregulated native and foreign genes for candidates with known roles in immunity in other invertebrate models^60,98,99^, such as Toll-like receptor genes (TLRs), caspases, thioester proteins, scavenger receptors, Gram-negative bacterial proteins (GNBPs) and peptidoglycan recognition proteins (PGRPs). We found little evidence for major upregulation of these gene families during exposure to the pathogen, though many of these pathways are detectable in bdelloid genomes. For example, for TLRs, there are 120 and 133 proteins with significant matches to the Toll/interleukin-1 receptor-like (TIR) domains (PF01582 and PF13676) in *A. vaga* and *A. ricciae*, respectively. However, we found only three putative *A. vaga* TLR genes to be significantly DE on exposure to the pathogen, and these were downregulated at T24; nine were found in *A. ricciae*, all but one downregulated. These observations do not exclude a role in immunity, since TLRs act as signalling components in other invertebrates^99,100^, and may be expressed constitutively rather than upregulated in response to pathogens. For caspases, we detected 101 and 70 proteins with a significant match to the Peptidase_C14 (PF00656) domain in *A. vaga* and *A. ricciae* respectively. Many of these were marked as HGT_C_ from bacteria (i.e., putative metacaspases) but in fact are highly divergent from any known caspase homologs found in UniProt (**Fig. 3f**), and their origin and evolution remain obscure. Similar to TLRs, only a relatively small number of putative caspases were found to be significantly differentially expressed on exposure to the pathogen, and the majority were downregulated. One possibility is that some genes with a role in immunity were downregulated in our experiment because they respond to non-fungal pathogens, such as bacteria. In nematodes, induction of antifungal defences is correlated with selective repression of antibacterial immune response genes^101^, perhaps to focus resources on an immediate threat or balance biochemical trade-offs in defence mechanisms^102^. Functional inferences about these genes are complicated by the substantial phylogenetic distance of rotifers from well-characterised models of invertebrate immunity^60,103^, as well as putative horizontal origins in some cases.

## Conclusions

We find that genes acquired by bdelloid rotifers from non-metazoan sources play a disproportionate role in a specific transcriptional response to infection by a fungal pathogen. The speed and scale of this response was greater in the more resistant of the two host species we compared, though expression differences were subtle relative to a fourfold difference in susceptibility. The enrichment of foreign genes was twice as pronounced in the response to pathogens as compared with desiccation, suggesting that their expression patterns are specific to biotic defence rather than a general reaction to stress or physical damage. Since virtually nothing is currently known about the genetics of immunity in rotifers, and tools for assessing the functions of candidate genes in bdelloids are lacking, the precise roles of the differentially regulated genes remain to be determined. Nonetheless, we hypothesise that RNA ligase and glucanase functions acquired from bacteria and other sources have been co-opted for immune defence against fungi in particular. While some putative horizontally acquired genes might in fact represent novel gene families with a deep or obscure metazoan heritage, the horizontal origin of the genes we highlight is well supported by phylogenetic evidence grouping them closely with non-metazoan orthologs.

Overall, our findings are consistent with the hypothesis that horizontal gene transfer contributes to bdelloid defence against pathogens. This could be interpreted to support the view that diseases pose a particular challenge for asexual lineages^13,14^, with special measures required for longstanding parthenogenetic lineages to keep up^9,27,28^. But to what extent would this process help bdelloids overcome the presumed deficiencies of long-term asexuality in combatting disease? Many of the genes that were upregulated in infected populations were acquired prior to the common ancestor of *A. vaga* and *A. ricciae*, and some appear to be shared even more deeply among bdelloid rotifers. Such genes could provide an enhanced biochemical repertoire for combatting pathogens overall, potentially including functions unavailable to other animals, but they would not create variation in bdelloid populations at the speed or scale theoretically required to sustain rapid coevolution^15,104–108^. Nonetheless, a handful of upregulated foreign genes were restricted to just one *Adineta* species and therefore interpreted as more recently acquired. It is possible that new genes do arrive at a fast enough rate to diversify immune function at the scale of lineages and species, and thereby provide some variation to counter long-term extinction, especially if ecological factors^56,58^ help susceptible rotifer clones avoid coevolving pathogens in the shorter term.

## Materials and methods

### Rotifer and pathogen isolates

Animals belonging to the species *Adineta ricciae*^80^ and *A. vaga*^109,110^ were isolated respectively in 1998 from mud in Australia^80^ and ca. 1984 from moss in Italy^111^. These have been propagated clonally in continuous long-term culture across several laboratories^43,48,112,113^ and annotated genome assemblies are available for both lines^43,46^. Our cultures were maintained in 60mm plastic Petri dishes in sterilised distilled water, fed with *Escherichia coli* (OP50) and *Saccharomyces cerevisiae* (S288c) and subcultured approximately once per month. They were stored at 20°C in an illuminated incubator (LMS, Kent, UK) with a 12:12 hour light:dark cycle.

The fungal pathogen *Rotiferopthora globospora*^79^ was found attacking co-occurring rotifers of the genus *Adineta* in soil in northern New York^57^. A pure culture on potato dextrose agar (PDA) was obtained in December 2008 using methods described elsewhere^114^, and deposited in April 2009 with the USDA Agricultural Research Service Collection of Entomopathogenic Fungal Cultures (ARSEF) for long-term cryogenic storage under the accession code ARSEF 8995. Frozen mycelium was retrieved and revived from this collection in 2013, and since maintained in serial subculture on PDA at 20°C with a 12:12 hour light:dark cycle and with transfers every 4 months. This pathogen strain has no recent history of laboratory co-passage with either of the two rotifer hosts tested here and all three were isolated on different continents. However, *R. globospora* appears to be globally distributed—it was originally described from New Zealand^79^ and has also been recorded in Japan (CGW, pers. obs.), in both cases attacking *Adineta*. Therefore, either or both host species could plausibly have encountered this pathogen in nature, especially given the high dispersal capacity and global distribution of bdelloid rotifers^58,115–117^.

Infection by *R. globospora* is initiated when a host ingests infectious spores (conidia), which are spherical and approximately 3.5μm in diameter^79^. These lodge in the mouth or oesophagus, at which point the animal stops feeding and contracts within a few minutes (**Fig. 1a**). Over the next 6–8 hours, the conidium produces a thin germ tube that penetrates the gut wall and swells into assimilative hyphae, which begin to invade and digest the surrounding host tissue after about 12–24 hours, as the infection becomes established. By 36–48 hours, fungal hyphae have filled most of the body cavity and the host is dead. Between 48–72 hours, hyphae begin to emerge through the host integument, and will eventually differentiate to form two types of spores: a handful of thick-walled resting spores, and hundreds of fresh infectious conidia (**Fig. 1a**).

Because *Rotiferophthora* can only complete its life cycle by killing its host, these fungi are inherently highly virulent pathogens, and have been described as “devastating” to populations of *Adineta*^79^. *R. globospora* (ARSEF 8995) has been shown to exterminate laboratory populations of a sympatric *Adineta* c.f. *vaga* clone within 28 days of initial exposure to a low density of conidia^57^. However, even if rotifer individuals are inevitably killed once the pathogen becomes established, the possibility remains that an individual host can resist the initial attack and prevent an ingested spore from establishing a successful infection, or delay its progression. Even partial resistance at an early stage could dramatically slow the spread of an epidemic^118^, giving clonal relatives time to escape the infested habitat^56,58^ before the local population is exterminated. Early-acting mechanisms of resistance to *Rotiferophthora* ought therefore to be favoured in bdelloids, corresponding to the innate immune pathways identified in other invertebrates facing virulent fungal pathogens^101,119–121^.

### Infection assays

To quantify resistance to *R. globospora* under standard conditions, rotifers were transferred by pipette from stock populations to a 2mL droplet of sterile distilled water, where eggs, corpses and food from the cultures were washed away. Adult individuals were transferred to 96-well plates (Thermo-Fisher), with approximately 11 animals per well (mean: 11.0, SD: 3.5) in 60μL of sterilised, distilled water. Rotifers were counted using a compound microscope (Nikon Eclipse E400), noting whether each animal was active (feeding or locomoting, **Fig. 1a**), contracted (with head withdrawn) or dead. Animals that died during the transfer (<1.5% of the total) were physically removed where possible, or recorded so they could be excluded from later counts.

To obtain pure suspensions of conidia, approximately 5×5mm of freshly subcultured sporulating mycelium on PDA was moved to 500μL of sterile distilled water in a 1.5mL Eppendorf tube. After vortexing, 400μL of conidial suspension was transferred to a fresh sterile tube and conidial density was measured using a haemocytometer and lactophenol cotton blue stain. An inactivated inoculum was prepared simultaneously as a control, by exposing an aliquot of conidia in water to a germicidal ultraviolet lamp (25W, 253.7nm) at a distance of 10cm for 45 minutes, with regular vortexing to ensure all spores were irradiated. This treatment was successful, as none of the rotifers in control wells treated with irradiated spores became infected.

Wells were inoculated with 8μL of freshly prepared conidial suspension at a density of 125 spores μL^-1^. Negative control wells received 8μL of distilled water or inactivated spore suspension. The final density of spores in each well was high (ca. 15 conidia μL^-1^) to ensure every animal was exposed to the pathogen in a synchronised pulse. This appeared to work, because 94% of animals in experimental wells were contracted 8 hours after exposure, versus a baseline of 3% of animals contracted in control wells. Plates were then stored in incubators (LMS 300NP) at 20°C, with a 12:12 hour light:dark cycle.

After 48 and 72 hours, rotifers were counted again, and classed as active, contracted, killed by infection or otherwise dead. A rotifer was considered to be killed by infection if at least one hypha had emerged through the integument from the interior. This criterion was unambiguous, in contrast with the difficulty of determining whether a fungal infection had established inside a contracted animal. By 72h, the proportion of infected animals in experimental wells appeared to have stabilised. In a subset of wells recounted at 96h, only a small fraction of animals (~6.5%) had newly developed visible infections. The relative risk of infection for *A. vaga* versus *A. ricciae* at 96h (RR 2.94, 95% CI 1.93–4.46; *Z* = 5.06, *P* < 0.0001) had not narrowed significantly since 72h (ratio of RR 0.78, 95% CI 0.47–1.31, *Z* = −0.926, *P* = 0.354), but the survivors had reproduced, making further tracking of the originally exposed cohort difficult. We therefore took the 72h timepoint as the standard measure of infection mortality. Individuals in negative control wells never became infected, whether they received water or sterilized spores, and the background death rate was ~2%.

Inoculation trials with both *A. ricciae* and *A. vaga* were replicated on multiple occasions using at least three separate source dishes for each species prepared at different times (in some cases different years), and separately prepared cultures and inocula of fungi, with each set of trials replicated in at least three wells with accompanying negative controls in appropriately blocked designs. The total number of animals exposed across all trials was 216 for *A. ricciae* and 189 for *A. vaga* (**Fig. 1b**). Rates of mortality from *R. globospora* infection were consistent across these trials, as was the dramatic difference in susceptibility between the species (**Fig. 1b**). One set of well trials was run simultaneously and in close linkage with the RNA-seq experiment described below, using leftover rotifers from the same source populations, and inoculating wells with the same live and irradiated pathogen suspensions at the same time. This enabled us to infer the timings and presumed final outcomes of infections in the RNA tubes, even though those animals could not be directly counted and were sacrificed for RNA by 24h. The results of the RNA-linked well experiments were similar to earlier trials: mean infection mortality at 72h was 18% for *A. ricciae* and 79% for *A. vaga*.

### RNA-seq experimental design

Rotifers for the RNA-seq experiment were reared in eight replicate Petri dishes per species, with ca. 50 founders per dish, all from the same clonal laboratory line of each species. These were fed only with *E. coli* (OP50, 5 x 10^8^ cells per dish) in distilled water. *S. cerevisiae* was omitted to avoid introducing further eukaryotic transcripts, and because gene expression by rotifers metabolising a fungal food source might complicate inferences about transcription in response to a fungal pathogen. The RNA-linked well assays described above yielded similar results to earlier trials, so the omission of *S. cerevisiae* as a food did not affect infection outcomes. When the *A. ricciae* dishes were initially established, each was inoculated with 50μL of 5μm-filtered water from the *A. vaga* source population, and vice-versa, to homogenise any co-cultured bacterial communities whose composition might differ between the cultured lines of the two species, and limit this as a potential source of gene expression differences. Dishes of the two species were stored in evenly interspersed blocks while the populations were growing.

Rotifers were counted and harvested after 4 weeks, when the mean population size was 2965 per dish for *A. ricciae* and 1688 for *A. vaga*, which reproduces more slowly^85^. Each dish was gently washed, using several water changes to float and pour off eggs, bacterial cells and corpses, while active animals adhered to the plastic. Cleaned rotifers were detached from the plastic by dropping distilled water onto them from a 10mL syringe at 40cm height, followed by repeated aspiration and forcible expulsion of medium using a P1000 pipetter and a 1mL tip. Physical detachment was preferred over a salt or cold shock approach^43^, to reduce the chance of inducing transcriptomic, immunological or behavioural changes. Suspended rotifers were poured into a 50mL centrifuge tube and pelleted at 3000 x g for 5 minutes using a swing-bucket cradle. All but 1.5mL of supernatant was removed, then a vortex shaker was used to resuspend the rotifer pellets and 1mL was transferred immediately by pipette to a 1.5mL Eppendorf tube. This process was repeated to yield 16 replicate 1.5mL tubes for each species. Each tube was centrifuged at 17000 x g for 3 minutes, and all but 100μL of water was removed from the rotifer pellet. The pelleted rotifers were left overnight to recover and redistribute themselves around the submerged interior of the tube. We estimate that the efficiency of rotifer recovery via this method is about 70%, based on numbers of animals left over in plates or tubes, giving approximately 1000 animals per tube for *A. ricciae* and 600 for *A. vaga*.

The 16 tubes for each species were randomly allocated to receive either live or irradiated pathogen spores, and to have RNA extracted either 7 or 24 hours later, with each combination replicated four times. Irradiated spore suspensions, as described above, were used as a control treatment to account for physical, chemical, or nutritional effects of ingesting fungal cells, so that all else was equal except for pathogenic activity. RNA sampling times were chosen to correspond to the early stages of infection^114^—by 7h, germ tubes from ingested spores have attempted to infiltrate the host, but assimilative hyphae would not yet have become established. By 24h, assimilative hyphae would be present if the germ tube had been successful, but these would not yet have colonised the host extensively or killed it. Each tube was inoculated with 20μL of live or irradiated spore suspension at a density of 500 spores μL^-1^, for a total of 10,000 spores and a final density of 80 conidia μL^-1^, to ensure every animal was exposed as synchronously as possible. Tubes were incubated upright at 20°C in a blocked layout until RNA extraction.

### RNA extraction and sequencing

Total RNA was extracted from each tube at the appropriate timepoint using an RNeasy Mini kit (Qiagen #74104), following the manufacturer’s protocol for animal tissues. To lyse and homogenise the rotifers, tubes were centrifuged at 17,000 x g for 3 minutes and 100μL of excess water was removed, leaving rotifers pelleted in approximately 20μL of water. After adding 50μL of Buffer RLT, the pellet was immediately disrupted and homogenised for 1 minute by pulsed application of a pellet pestle attached to a cordless motor (Kimble). A further 250μL of Buffer RLT was added to stabilise the lysate and rinse residue from the pestle, before proceeding with the manufacturer’s protocol, including an on-column DNase I digestion step. Lysis and stabilisation were completed for all tubes within 30 minutes of the target timepoint, in a balanced order with respect to treatment group and species.

RNA was eluted in 32μL of RNase-free water and 1.5μL aliquots were analysed using a Nanodrop 2000 (ThermoFisher). Spectrophotometric measurements were used to select the three replicates with the highest RNA concentrations from each treatment group for further analysis and sequencing, so that downstream steps had three biological replicates. These 24 tubes were frozen at −80°C and shipped on dry ice to the University of Edinburgh (Edinburgh, UK), where further quality control was undertaken by Edinburgh Genomics, including RNA quantitation with a Qubit 2.0 fluorometer in duplicate for each sample, and RNA ScreenTape analysis with an Agilent 2200 TapeStation system to assess RNA integrity. All 24 samples proceeded to cDNA library preparation, using the TruSeq stranded mRNA kit (Illumina) to enrich for polyadenylated transcripts. The indexed libraries were sequenced in multiplex on an Illumina NovaSeq 6000 at Edinburgh Genomics, using an SP flow cell to generate 50-base paired end read libraries with 200 bp inserts.

A total of 102.9 Gb of raw sequencing data were generated over the 24 RNA libraries. After data filtering and quality-control, 78.5 million reads were retained per library (94.2 Gb total data, Supplementary Table 1). Over 99% of filtered reads mapped to the *A. vaga* and *A. ricciae* reference genomes (Supplementary Table 2). Filtered data were *de novo* assembled using the Trinity software v2.8.9^122,123^, resulting in 118,860 and 63,749 transcripts representing 41,527 and 35,079 ‘genes’ for *A. ricciae* and *A. vaga*, respectively (Supplementary Table 2). Transcriptomes showed G + C proportions that were consistent with their respective genomes (*A. ricciae* 37.6%; *A. vaga* 33.0%) and high ‘completeness’ scores as measured by BUSCO analysis (>97% of core eukaryote genes completely recovered in both cases). Transcript-to-genome mapping rates were >99% in both species, with 87.1% (*A. ricciae)* and 81.8% (*A. vaga*) of introns correctly called based on previously published gene models^46^.

### Data filtering and quality control

Raw RNA-seq sequencing reads were quality and adapter trimmed using BBTools ‘bbduk’ v38.73 with the parameters ‘ktrim=r k=23 mink=11 hdist=1 tpe tbo’, and error-corrected using BBTools ‘tadpole’ with the parameters ‘mode=correct tossjunk tossuncorrectable’ (https://sourceforge.net/projects/bbmap/). Unwanted reads derived from ribosomal RNA (rRNA) were removed by mapping the data to the SILVA rRNA database^124^ using BBTools ‘bbmap’ with the parameters ‘local=t outu=filtered_R#.fq.gz’ (i.e. retaining only unmapped pairs of reads). Sequences derived from fungal contaminants were also removed using the same approach, mapping to sequenced genomes of fungi in the family *Clavicipitaceae* (NCBI:txid34397). The proportion of reads mapping to the *A. ricciae* or *A. vaga* target genome was assessed using STAR v2.7.3a^125^ with the parameter ‘--twoPassMode Basic’. Final data quality was assessed visually using FastQC^126^ and MultiQC^127^. All raw sequencing data have been deposited in the relevant International Nucleotide Sequence Database Collaboration (INSDC) database with the Study ID PRJEB39927 (see Supplementary Table 1 for run accessions).

### Differential expression analysis

Transcript quantification was performed using Salmon ‘quant’ v0.14.1^128^, using the gene annotations of Nowell et al.^46^ as the target transcriptomes. Short transcripts <150 bases were removed prior to analysis, and genomic scaffolds were appended to each transcriptome data as ‘decoys’ prior to quantification as recommended in the Salmon documentation. Relationships between biological replicates within and between samples were checked visually using utility scripts from the Trinity software^122,123^. Statistical analysis of the resulting count matrix was performed with DESeq2^129^, which uses negative binomial generalized linear models to test for differential expression. *P*-values were adjusted for multiple testing using the Benjamini-Hochberg method^130^ to control the false discovery rate (FDR). Stringent thresholds of FDR < 1e–3 and log_2_ fold-change > 2 (i.e. 4-fold difference in expression) were used to define an initial set of differentially expressed genes for downstream analysis.

To assess the quality of the *A. vaga* and *A. ricciae* gene predictions^46^, a transcriptome was assembled *de novo* from the filtered RNA-seq data using Trinity v2.8.9^122^ with default parameters, and mapped to the genomes of *A. ricciae*^46^ and *A. vaga*^43^ using Minimap2^131^ v2.17 with the parameters ‘-ax splice -C5’. The degree of correspondence between the predicted gene models and the assembled transcriptome was then assessed using ‘paftools.js junceval’, which provides a count of the number of correctly predicted introns based on the exact agreement between the location of the intron-exon junction in the annotation file and gaps in the transcriptome alignments. Gene completeness of assembled transcriptomes was also measured by Benchmarking Universal Single-Copy Orthologs (BUSCO) analysis^132^, using the core eukaryote and metazoa gene sets (*n* = 303 and 978 query genes, respectively).

### Functional annotation

Protein sequences from the *A. ricciae* and *A. vaga* reference genomes were aligned to the SwissProt^133^ and Pfam^134^ sequence databases using BLAST^135^ and HMMER v3.3 (http://hmmer.org/) respectively. Signal peptide cleavage sites and transmembrane helices were identified using SignalP v4.1^136^ and TMHMM v2.0^137^ respectively. Protein domains were further identified using InterProScan v5^138^. Functional annotations were assimilated using the Trinotate v3.2.0^139^ pipeline.

### Horizontal gene transfer detection

Putative HGT candidate genes (HGT_C_) were determined from the recent analysis of Jaron et al.^70^. Briefly, proteins were aligned to the UniProt90 sequence database^133^ using Diamond ‘blastp’ v0.9.21^140^ with the parameters ‘--sensitive -k 500 -e 1e-10’. For each protein, a HGT ‘index’ (*h*_U_) was computed based on the alignment bitscores to the best-matching hits from the Metazoa (*B*_IN_) and non-Metazoa (*B*_OUT_) with the formula: *h*_U_ = *B*_OUT_ – *B*_IN_^68^. A ‘consensus hit support’ (CHS) score was also calculated as the proportion of secondary hits in agreement with the result based on the *h*_U_^46,141^. Proteins were classified as HGT_C_ if *h*_U_ > 30, CHS > 90% and were physically linked (i.e., found on the same genomic scaffold) to a gene of unambiguous metazoan origin (*h*_U_ < 0). Phylogenetic trees of selected HGT_C_ and their putative co-orthologs from the UniRef90 database were constructed using IQ-TREE v1.6.12 with the parameters ‘-alrt 1000 -bb 5000 -m TEST’^142–144^.

### Gene orthology

Orthologous relationships between *A. ricciae* and *A. vaga* genes was determined using OrthoFinder v2.3.12^145^ with default parameters. Proteomes from nine other rotifer species^46,47,146^ were included in the analysis to aid orthology inference.

## Acknowledgements

Genome sequencing was performed by the UK Natural Environment Research Council (NERC) Biomolecular Analysis Facility at Edinburgh Genomics at the University of Edinburgh (NBAF-Edinburgh). The authors wish to thank Matthew Arno, Colin Sharp, Isobel Eyres, Chiara Boschetti, Rebecca Allen, and Mariya P. Dobreva.

## Supplementary materials

**Supplementary Table 1.**
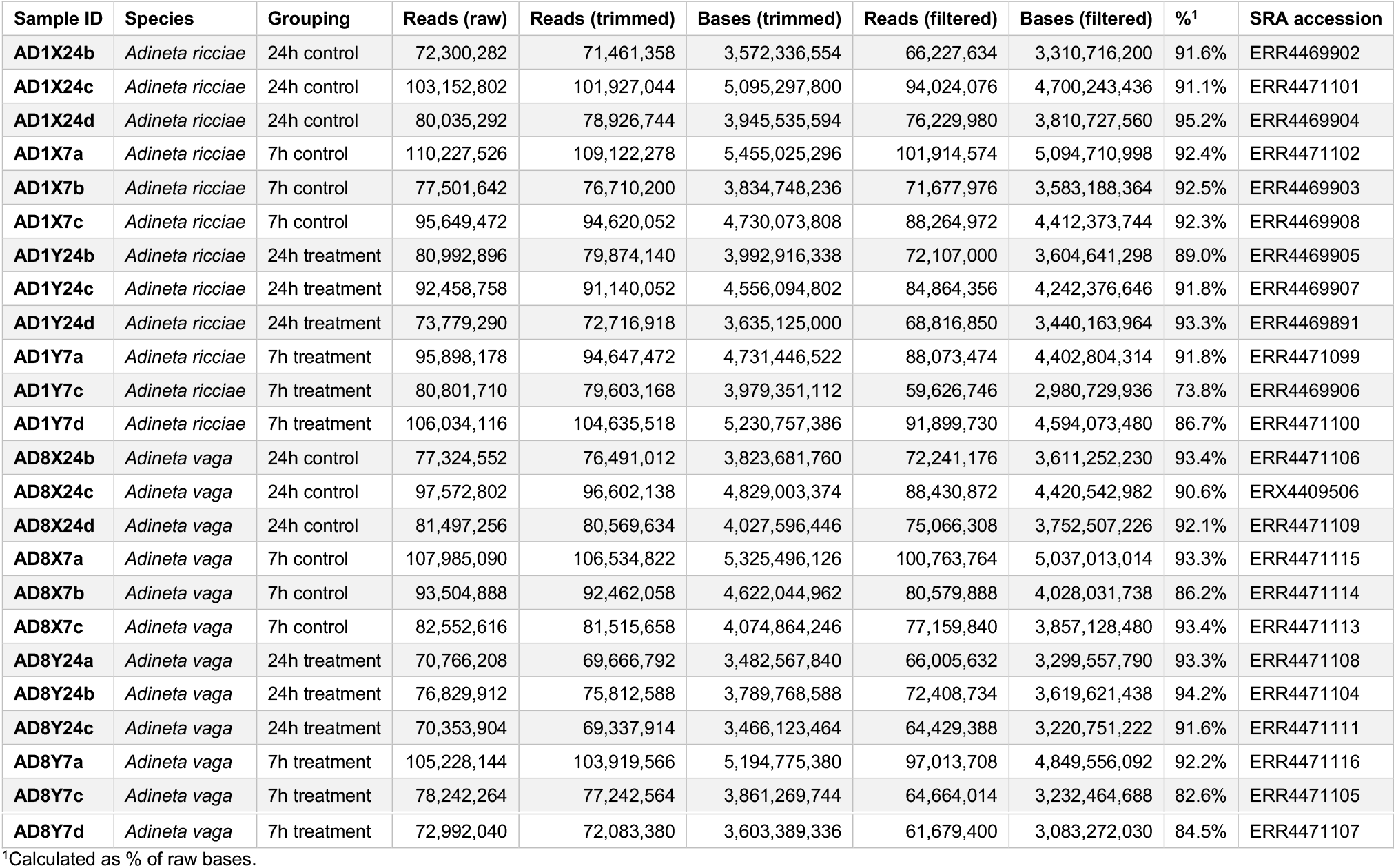
Data counts for raw and filtered sequencing data.

**Supplementary Table 2.** Assembly and quality metrics for Trinity *de novo* transcriptomes.

|  | *Adineta ricciae* | *Adineta vaga* |
| --- | --- | --- |
| **Number of transcripts** | 118,860 | 63,749 |
| **Mean** | 1,400 | 1,396 |
| **Median** | 1,157 | 1,080 |
| **Max** | 31,021 | 20,120 |
| **Total span** | 166,388,199 | 88,983,009 |
| **GC%** | 37.6 | 33.0 |
| **Number of 'genes'** | 41,527 | 35,079 |
| **Total span of 'genes'** | 48,809,330 | 41,442,308 |
| **BUSCO<sup>1</sup> eukaryote (*n* = 303)** | C:97.7%[S:10.9%,D:86.8%],F:1.7% | C:99.0%[S:57.4%,D:41.6%],F:0.7% |
| **BUSCO metazoa (*n* = 978)** | C:93.4%[S:11.5%,D:81.9%],F:1.4% | C:93.8%[S:51.8%,D:42.0%],F:1.0% |
| **Mapped reads to transcriptome** | 99.4% | 99.5% |
| **Mapped transcripts to genome<sup>2</sup>** | 118,277 (99.5%) | 63,379 (99.4%) |
| **Introns predicted from GFF<sup>3</sup>** | 478,936 | 276,708 |
| **Correct introns from mapping** | 417,174 (87.1%) | 226,332 (81.8%) |

<sup>1</sup>BUSCO codes: **C**, complete; **S**, complete and single copy; **D**, complete and duplicated; **F**, fragmented
<sup>2</sup>Genome accessions *A. ricciae*: GCA\_900240375.1; *A. vaga*: GCA\_000513175.1<sup>43,46</sup>.
<sup>3</sup>GFF annotations from Nowell et al.<sup>46</sup>.

**Supplementary Table 3.** Pearson’s product-moment correlations in log_2_ fold-chanαe expression.

| Test<sup>1</sup> | Species | Timepoint<sup>2</sup> | DE type<sup>3</sup> | Pearson's *R* (95%-CI) | *P*-value |
| --- | --- | --- | --- | --- | --- |
| Timepoints | *A. vaga* | Both | Up | 0.84 (0.84–0.85) | <0.001 |
|  |  |  | Down | 0.67 (0.66–0.68) | <0.001 |
|  | *A. ricciae* | Both | Up | 0.82 (0.82–0.83) | <0.001 |
|  |  |  | Down | 0.66 (0.65–0.67) | <0.001 |
| Within genomes | *A. vaga* | T7 | Up | 0.43 (0.42–0.44) | <0.001 |
|  |  |  | Down | 0.50 (0.49–0.52) | <0.001 |
|  |  | T24 | Up | 0.43 (0.42–0.44) | <0.001 |
|  |  |  | Down | 0.50 (0.49–0.51) | <0.001 |
|  | *A. ricciae* | T7 | Up | 0.49 (0.48–0.50) | <0.001 |
|  |  |  | Down | 0.39 (0.38–0.40) | <0.001 |
|  |  | T24 | Up | 0.60 (0.60–0.61) | <0.001 |
|  |  |  | Down | 0.42 (0.41–0.43) | <0.001 |
| Between genomes | Both | T7 | Up | 0.35 (0.34–0.36) | <0.001 |
|  |  |  | Down | 0.26 (0.25–0.27) | <0.001 |
|  |  | T24 | Up | 0.37 (0.36–0.37) | <0.001 |
|  |  |  | Down | 0.40 (0.39–0.40) | <0.001 |

<sup>1</sup>Explanation of tests: “Timepoints”, correlation in differential expression (DE) in the same gene across timepoints T7 and T24 (Supplementary Fig. 1); “Within genomes”, correlation in DE between inferred gene copies within genomes (including putative homologs, homoeologs and other gene copies) (Supplementary Fig. 2a–d); “Between genomes”; correlation in DE between inferred orthologs across species (Supplementary Fig. 2e–f). <sup>2</sup>Timepoints: “T7”, 7h post-infection; “T24”, 24h post-infection. <sup>3</sup>DE type: “Up”, genes that are upregulated in the treatment group (i.e., positive DE); “Down”, genes that are downregulated in the treatment group (negative DE). Note that all genes are included in correlations regardless of significance.

**Supplementary Table 4.** Proportion of *A. vaga* DE genes shared between pathogen and desiccation conditions.

| Gene type | DE type |  | T7 | T24 | Ent | Rec |
| --- | --- | --- | --- | --- | --- | --- |
| All DE genes | Up | T7 | **452** | 89.6% | 7.1% | 21.9% |
| All DE genes | Up | T24 | 37.1% | **1093** | 9.3% | 20.9% |
| All DE genes | Up | Ent | 5.7% | 18.3% | **558** | 62.7% |
| All DE genes | Up | Rec | 5.5% | 12.6% | 19.4% | **1807** |
| All DE genes | Down | T7 | **89** | 66.3% | 2.2% | 0.0% |
| All DE genes | Down | T24 | 8.8% | **674** | 2.1% | 11.3% |
| All DE genes | Down | Ent | 0.6% | 3.9% | **361** | 26.3% |
| All DE genes | Down | Rec | 0.0% | 5.7% | 7.2% | **1322** |
| HGT<sub>C</sub> | Up | T7 | **136** | 89.7% | 8.8% | 22.8% |
| HGT<sub>C</sub> | Up | T24 | 45.4% | **269** | 10.8% | 22.7% |
| HGT<sub>C</sub> | Up | Ent | 16.4% | 39.7% | **73** | 57.5% |
| HGT<sub>C</sub> | Up | Rec | 12.1% | 23.8% | 16.4% | **256** |
| HGT<sub>C</sub> | Down | T7 | **25** | 80.0% | 4.0% | 0.0% |
| HGT<sub>C</sub> | Down | T24 | 12.2% | **164** | 1.8% | 14.6% |
| HGT<sub>C</sub> | Down | Ent | 2.9% | 8.8% | **34** | 41.2% |
| HGT<sub>C</sub> | Down | Rec | 0.0% | 9.5% | 5.5% | **253** |

Values in bold on diagonals indicate the total number of genes in category (e.g., 136 HGT<sub>C</sub> genes upregulated at T7). Percentages in each cell then indicate the proportion of shared genes relative to the row total (e.g., 89.7% of 136 T7 upregulated HGT<sub>C</sub> genes are shared with T24 upregulated set). Green and red shading is to highlight the relative size of the shared fraction for up- and downregulated subsets, respectively.

**Supplementary Figure 1.**
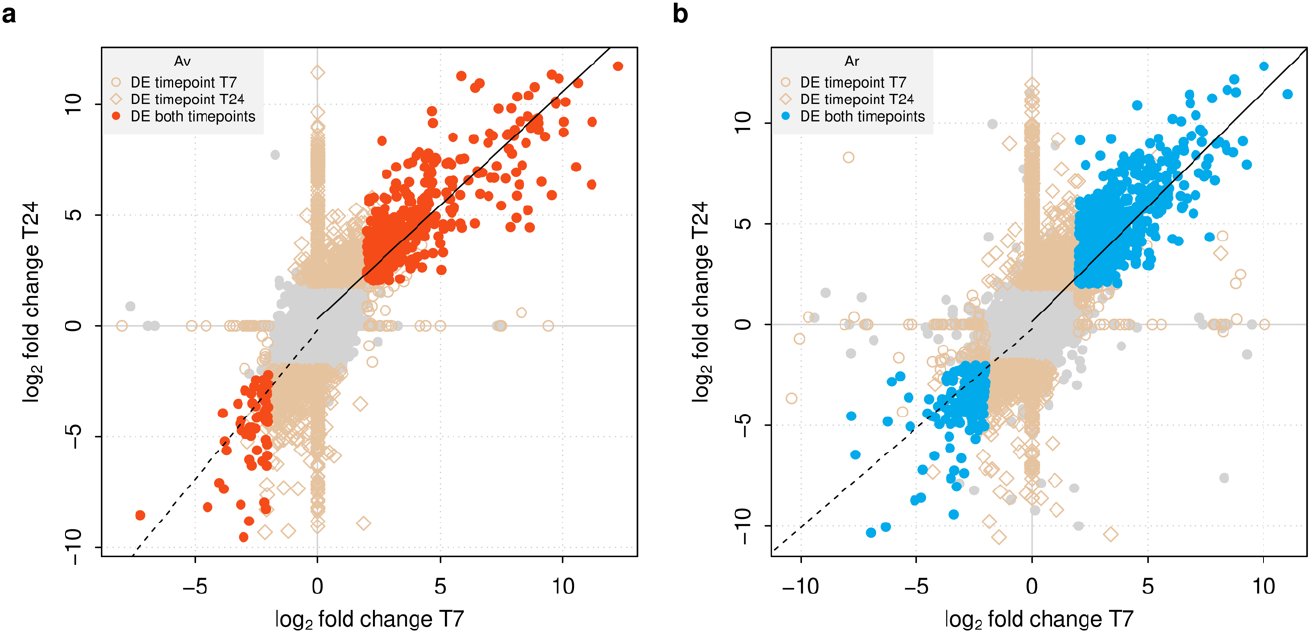
Correlation in DE between timepoints. Each point represents a single gene, and its log_2_ fold change in expression in treatment groups versus control groups at timepoint T7 (*X*-axis) versus timepoint T24 (*Y*-axis) for **a.** *A. vaga* and **b.** *A. ricciae*. Positive values represent upregulation in the treatment groups relative to control groups; negative values represent downregulation. Genes with significant DE in both timepoints are shown in red (*A. vaga*) and blue (*A. ricciae*); genes significant in one timepoint but not the other are shown with diamond and circle symbols (see legends). Genes with non-significant DE in both timepoints are plotted in grey. Solid black lines show the linear relationship for all upregulated genes (i.e., genes with log_2_ fold change > 0 in both timepoints; Pearson’s correlation *R* = 0.84 and 0.82 for *A. vaga* and *A. ricciae*, respectively; *P* < 2e–16 in both cases). Dashed black lines show the linear relationship for downregulated genes (log_2_ fold change < 0 in both timepoints; Pearson’s correlation *R* = 0.67 and 0.66 for *A. vaga* and *A. ricciae*, respectively; *P* < 2e–16 in both cases).

**Supplementary Figure 2.**
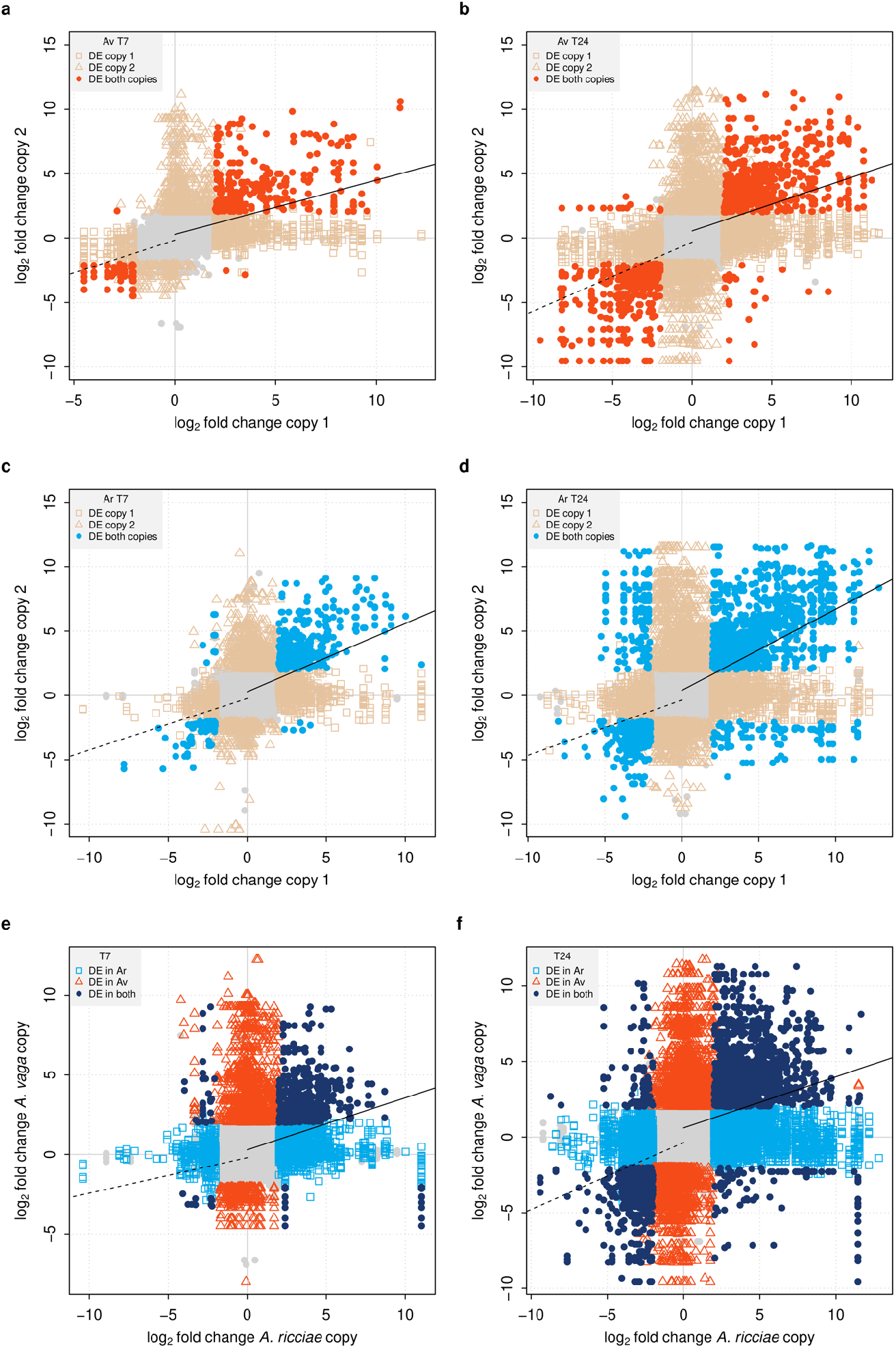
Correlation in DE between gene copies within and between genomes. Plots show the log_2_ fold change in expression in treatment groups versus control groups for a. gene copies within *A. vaga* at timepoint T7, b. gene copies within *A. vaga* at timepoint T24, c. gene copies within *A. ricciae* at T7, d. gene copies within *A. ricciae* at T24, e. gene copies between species at T7, and finally f. gene copies between species at T24. Note that each point represents a relationship between a pair of genes, not a gene itself (i.e., putative homologs, homoeologs or paralogs for within-genome comparisons, or orthologs for between-genome comparisons). Solid black lines show the linear relationship for all upregulated genes (i.e., genes with log_2_ fold change > 0 in both copies; Pearson’s correlation *R* = 0.43, 0.43, 0.49, 0.60, 0.35 and 0.37 for *A. vaga* and *A. ricciae*, timepoints T7 and T24 respectively; *P* < 2e–16 in all cases). Dashed black lines the linear relationship for downregulated genes (log_2_ fold change < 0 at both timepoints; Pearson’s correlation *R* = 0.50, 0.50, 0.39, 0.42, 0.26 and 0.40 for *A. vaga* and *A. ricciae*, timepoints T7 and T24 respectively; *P* < 2e–16 in all cases).

**Supplementary Figure 3.**
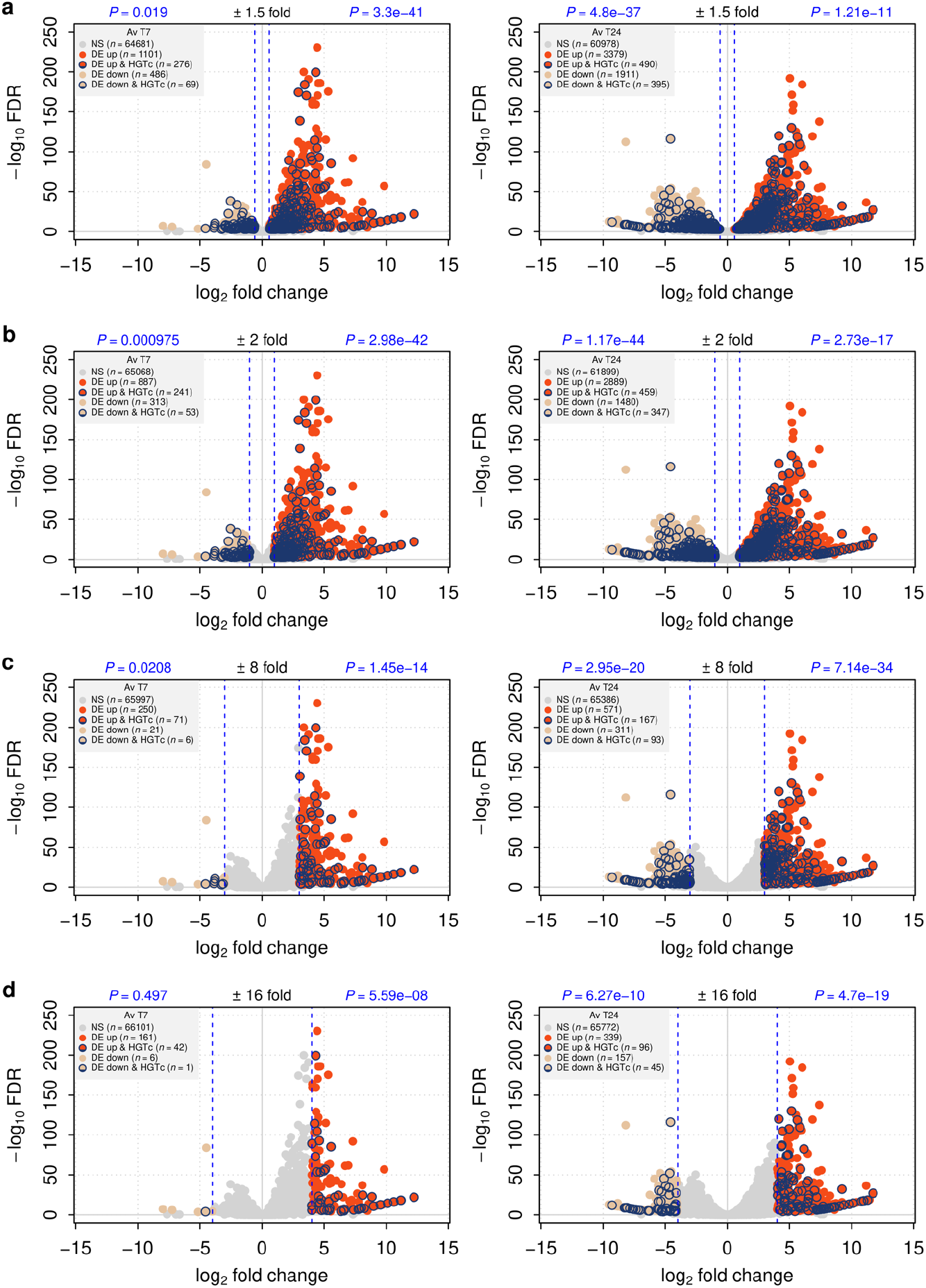
Effect of varying DE significance parameters on enrichment of HGT_C_ in *A. vaga*. Plots are arranged as previously. Dashed blue lines show ± log_2_ fold change equivalent to a 1.5-fold, b 2-fold, c 8-fold, and d 16-fold change in expression value to demark up- and downregulated subsets. Threshold for FDR < 1e–3 in all cases. *P*-values in blue show the probability of observing these data given the null hypothesis of no enrichment of HGT_C_ in corresponding subset (Fisher exact tests; see Methods).

**Supplementary Figure 4.**
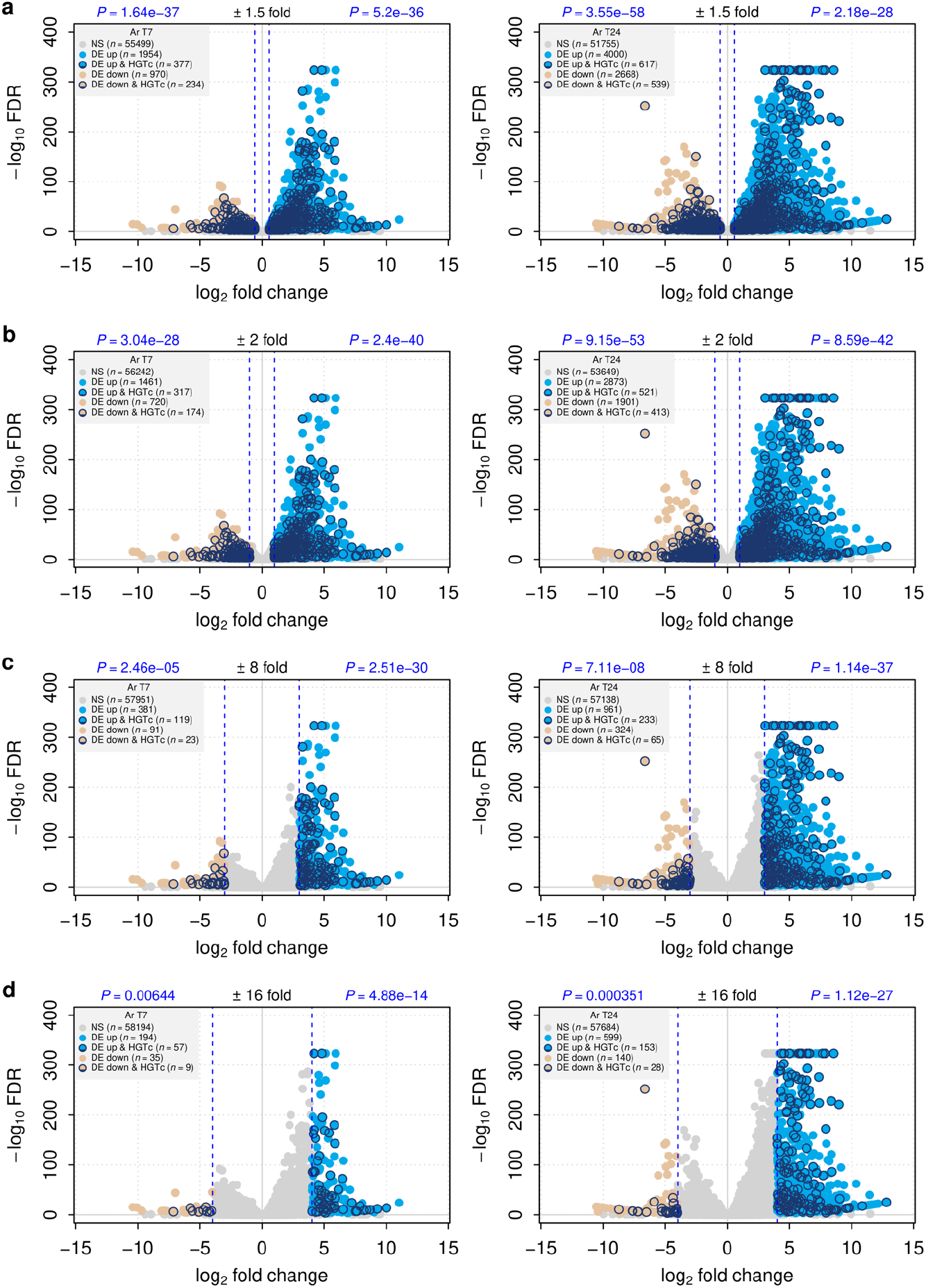
Effect of varying DE significance parameters on HGT_C_ enrichment in *A. ricciae*. Plots are arranged as previously. Dashed blue lines show ± log_2_ fold change equivalent to a 1.5-fold, b 2-fold, c 8-fold, and d 16-fold change in expression value to demark up- and downregulated subsets. Threshold for FDR < 1e–3 in all cases. *P*-values in blue show the probability of observing these data given the null hypothesis of no enrichment of HGT_C_ in corresponding subset (Fisher exact tests; see Methods).

**Supplementary Figure 5.**
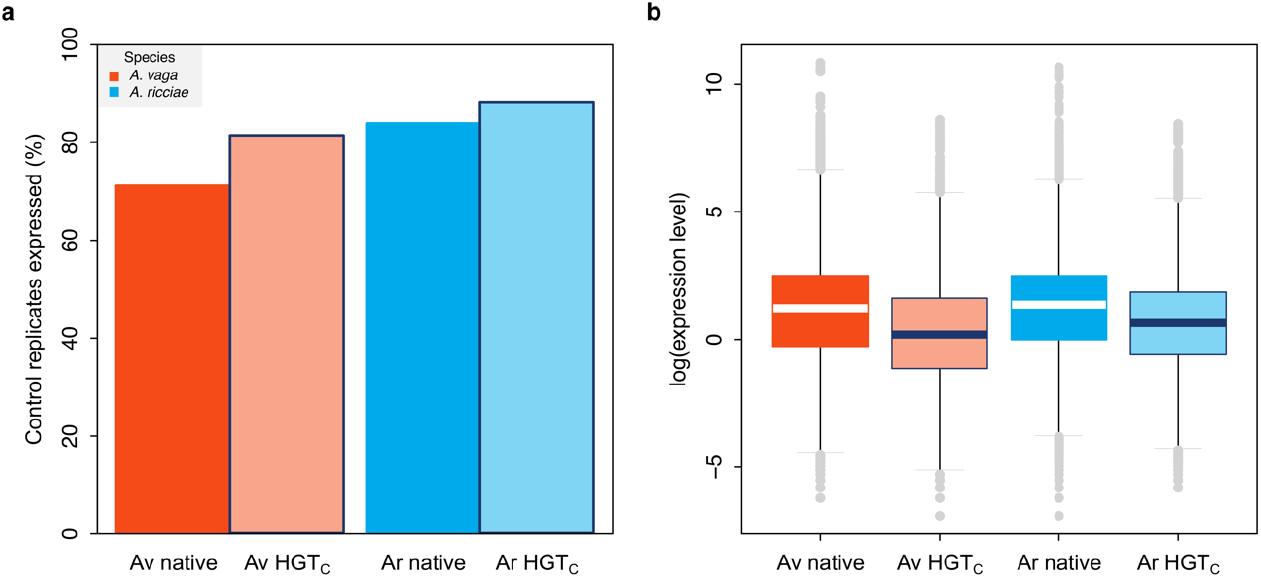
Expression of foreign genes in control groups. **a** Proportion of native (dark shade) and foreign (light shade with blue border) genes expressed in control groups (i.e., trimmed mean of M-values [TMM] > 0 in any control replicate). Fisher exact test for count data, for *A. vaga* odds ratio = 1.78 (95% CI = 1.73–1.82), for *A. ricciae* odds ratio = 1.45 (95% CI = 1.40–1.50); *P* < 0.001 in both cases. **b** Level of expression (log TMM) of native and foreign genes expressed in control replicates. Mean (log) TMM Av native = 3.10 ± 5.47 SD; Av HGT_C_ = 2.42 ± 4.57 SD; Welch two sample *t*-test, *t* = 74 (45,525 d.f.), *P* < 0.001; Mean (log) TMM Ar native = 2.98 ± 5.31 SD; Ar HGT_C_ = 2.61 ± 4.80 SD; Welch two sample *t*-test, *t* = 55 (38,831 d.f.), *P* < 0.001.

**Supplementary Figure 6.**
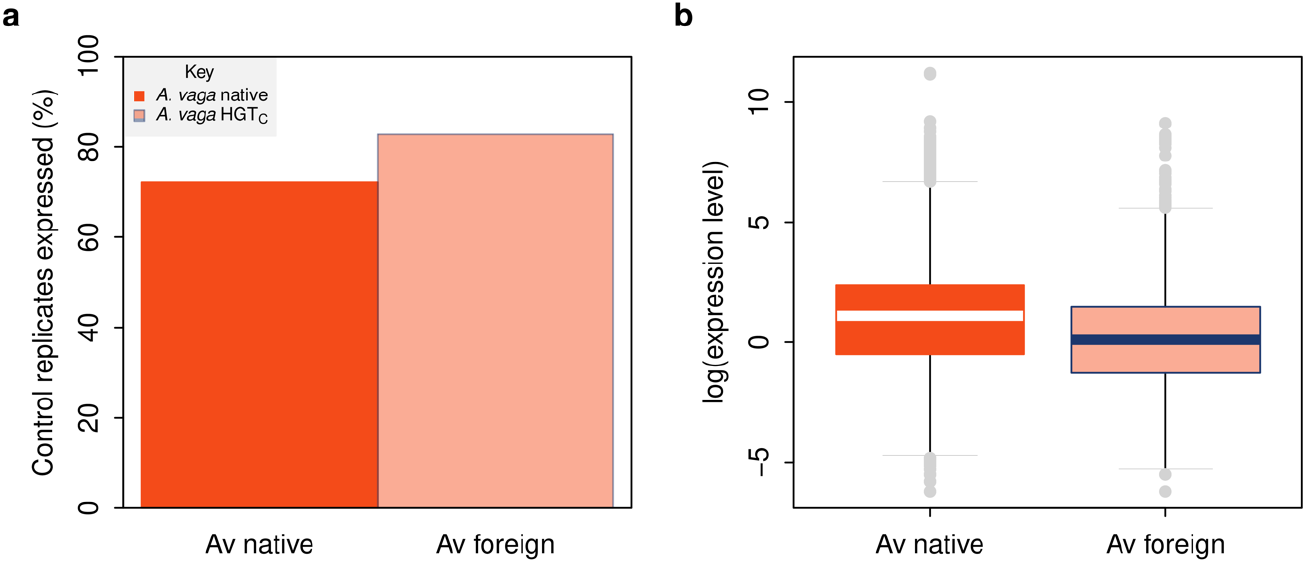
Gene expression in control replicates of *A. vaga* in response to drying. **a** Proportion of native (dark shade) and foreign (light shade with blue border) genes expressed in control groups (i.e., trimmed mean of M-values [TMM] > 0 in any control replicate). Fisher exact test for count data, odds ratio = 1.78 (95% CI = 1.77–1.91), *P* < 0.001. **b** Level of expression (log TMM) of native and foreign genes expressed in control replicates. Mean (log) TMM Av native = 3.05 ± 5.90 SD; Av foreign = 2.59 ± 5.26 SD; Welch two sample *t*-test, *t* = 52.2 (23,427 d.f.), *P* < 0.001.

**Supplementary Figure 7.**
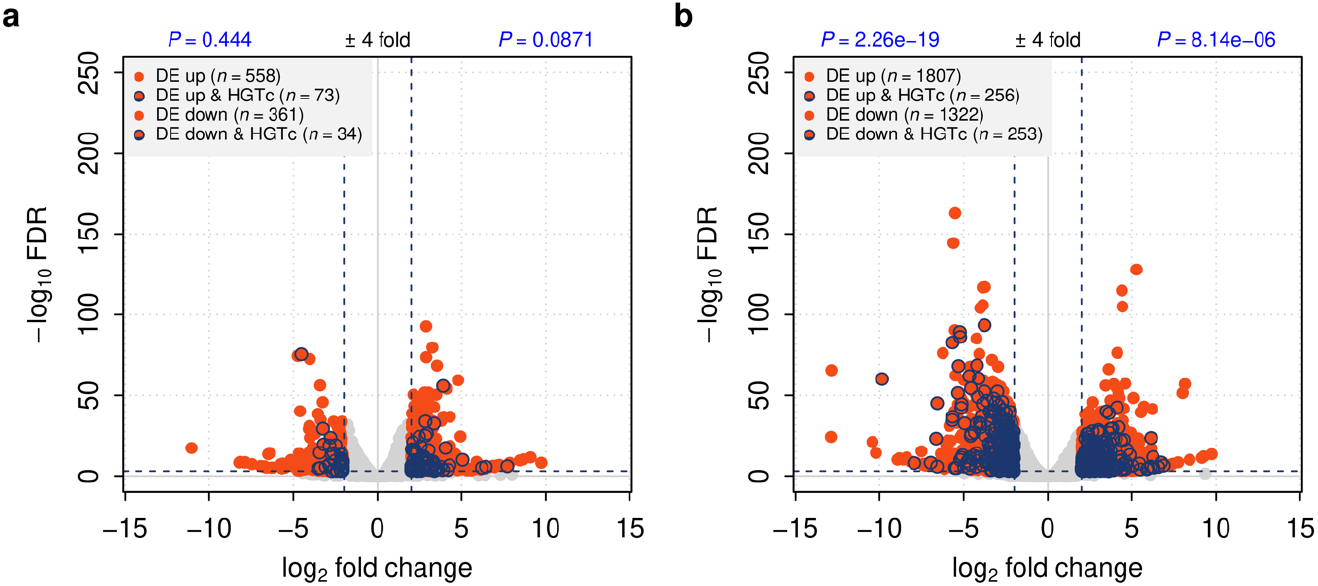
DE in *A. vaga* in response to drying. **a.** Animals entering desiccation; **b.** animals recovering from desiccation. Plots are arranged as previously. *P*-values in blue show the probability of observing these data given the null hypothesis of no enrichment of HGT_C_ in the corresponding subset (Fisher exact test; see Mmatethods).

## References

1. Darwin, C. On the two forms, or dimorphic condition, in the species of Primula, and on their remarkable sexual relations. Bot. J. Linn. Soc. 6, 77–96 (1862).

2. Williams, G. C. Sex and Evolution. 3–200 (Princeton University Press, Princeton, 1975).

3. Bell, G. The Masterpiece of Nature: The Evolution and Genetics of Sexuality. (University of California Press, Berkeley, 1982).

4. Maynard Smith, J. The Evolution of Sex. (Cambridge University Press, Cambridge, UK, 1978).

5. Gibson, A. K., Delph, L. F. & Lively, C. M. The two-fold cost of sex: Experimental evidence from a natural system. Evolution Letters 1, 6–15 (2017).

6. Lehtonen, J., Jennions, M. D. & Kokko, H. The many costs of sex. Trends Ecol. Evol. 27, 172–178 (2012).

7. Burt, A. Perspective: Sex, recombination, and the efficacy of selection—was Weismann right? Evolution 54, 337–351 (2000).

8. Speijer, D., Lukeš, J. & Eliáš, M. Sex is a ubiquitous, ancient, and inherent attribute of eukaryotic life. Proc. Natl. Acad. Sci. U. S. A. 112, 8827–8834 (2015).

9. Neiman, M., Meirmans, S. & Meirmans, P. G. What can asexual lineage age tell us about the maintenance of sex? Ann. N. Y. Acad. Sci. 1168, 185–200 (2009).

10. Moreira, M. O., Fonseca, C. & Rojas, D. Parthenogenesis is self-destructive for scaled reptiles. Biology Letters 17, 20210006 (2021).

11. Kondrashov, A. S. Classification of hypotheses on the advantage of amphimixis. J. Hered. 84, 372–387 (1993).

12. Meirmans, S. & Strand, R. Why are there so many theories for sex, and what do we do with them? Journal of Heredity 101 Suppl 1, S3–12 (2010).

13. Jaenike, J. A hypothesis to account for the maintenance of sex within populations. Evol. Theory 3, 191–194 (1978).

14. Hamilton, W. D. Sex versus Non-Sex versus Parasite. Oikos 35, 282–290 (1980).

15. Lively, C. M. A review of Red Queen models for the persistence of obligate sexual reproduction. J. Hered. 101 Suppl 1, S13–20 (2010).

16. Hamilton, W. D., Axelrod, R. & Tanese, R. Sexual reproduction as an adaptation to resist parasites (a review). Proc. Natl. Acad. Sci. U. S. A. 87, 3566–3573 (1990).

17. Agrawal, A. F. Similarity selection and the evolution of sex: revisiting the red queen. PLoS Biol. 4, e265 (2006).

18. Howard, R. S. & Lively, C. M. Parasitism, mutation accumulation and the maintenance of sex. Nature 367, 554–557 (1994).

19. West, S. A., Lively, C. M. & Read, A. F. A pluralist approach to sex and recombination. J. Evol. Biol. 12, 1003–1012 (1999).

20. Hodgson, E. E. & Otto, S. P. The red queen coupled with directional selection favours the evolution of sex. J. Evol. Biol. 25, 797–802 (2012).

21. Otto, S. P. & Nuismer, S. L. Species interactions and the evolution of sex. Science 304, 1018–1020 (2004).

22. Meirmans, S., Meirmans, P. G. & Kirkendall, L. R. The costs of sex: facing real-world complexities. Q. Rev. Biol. 87, 19–40 (2012).

23. Neiman, M., Meirmans, P. G., Schwander, T. & Meirmans, S. Sex in the wild: How and why field-based studies contribute to solving the problem of sex. Evolution (2018) doi:10.1111/evo.13485.

24. Gibson, A. K., Delph, L. F., Vergara, D. & Lively, C. M. Periodic, Parasite-Mediated Selection For and Against Sex. Am. Nat. 192, 537–551 (2018).

25. Tucker, A. E., Ackerman, M. S., Eads, B. D., Xu, S. & Lynch, M. Population-genomic insights into the evolutionary origin and fate of obligately asexual Daphnia pulex. Proc. Natl. Acad. Sci. U. S. A. 110, 15740–15745 (2013).

26. Greimann, E. S. et al. Phenotypic Variation in Mitochondria-Related Performance Traits Across New Zealand Snail Populations. Integr. Comp. Biol. 60, 275–287 (2020).

27. Judson, O. P. & Normark, B. B. Ancient asexual scandals. Trends Ecol. Evol. 11, 41–46 (1996).

28. Normark, B. B., Judson, O. P. & Moran, N. A. Genomic signatures of ancient asexual lineages. Biol. J. Linn. Soc. Lond. 79, 69–84 (2003).

29. Lively, C. M. & Morran, L. T. The ecology of sexual reproduction. Journal of Evolutionary Biology 27, 1292–1303 (2014).

30. Ejsmond, M. J. & Radwan, J. Red Queen Processes Drive Positive Selection on Major Histocompatibility Complex (MHC) Genes. PLoS Comput. Biol. 11, e1004627 (2015).

31. Lively, C. M. Coevolutionary Epidemiology: Disease Spread, Local Adaptation, and Sex. The American Naturalist 187, E77–82 (2016).

32. Ebert, D. & Fields, P. D. Host-parasite co-evolution and its genomic signature. Nat. Rev. Genet. 21, 754–768 (2020).

33. Donner, J. Ordnung Bdelloidea. 297 (Akademie Verlag, Berlin, 1965).

34. Segers, H. Annotated checklist of the rotifers (Phylum Rotifera), with notes on nomenclature, taxonomy and distribution. Zootaxa (2007).

35. Hubbs, C., Drewry, G. E. & Warburton, B. Occurrence and Morphology of a Phenotypic Male of a Gynogenetic Fish. Science 129, 1227–1229 (1959).

36. Palmer, S. C. & Norton, R. A. Taxonomic, geographic and seasonal distribution of thelytokous parthenogenesis in the Desmonomata (Acari: Oribatida). Exp. Appl. Acarol. 12, 67–81 (1991).

37. Smith, R. J., Kamiya, T. & Horne, D. J. Living males of the “ancient asexual” Darwinulidae (Ostracoda: Crustacea). Proc. Biol. Sci. 273, 1569–1578 (2006).

38. Soper, D. M., Neiman, M., Savytskyy, O. P., Zolan, M. E. & Lively, C. M. Spermatozoa Production by Triploid Males in the New Zealand Freshwater Snail Potamopyrgus antipodarum. Biological Journal of the Linnean Society 110, 227–234 (2013).

39. Fradin, H. et al. Genome Architecture and Evolution of a Unichromosomal Asexual Nematode. Curr. Biol. 27, 2928–2939.e6 (2017).

40. Boyer, L., Jabbour-Zahab, R., Mosna, M., Haag, C. R. & Lenormand, T. Not so clonal asexuals: Unraveling the secret sex life of Artemia parthenogenetica. Evol. Lett. (2021) doi:10.1002/evl3.216.

41. Arkhipova, I. & Meselson, M. Transposable elements in sexual and ancient asexual taxa. Proc. Natl. Acad. Sci. U. S. A. 97, 14473–14477 (2000).

42. Mark Welch, D. & Meselson, M. Evidence for the evolution of bdelloid rotifers without sexual reproduction or genetic exchange. Science 288, 1211–1215 (2000).

43. Flot, J.-F. et al. Genomic evidence for ameiotic evolution in the bdelloid rotifer *Adineta vaga*. Nature 500, 453–457 (2013).

44. Debortoli, N. et al. Genetic Exchange among Bdelloid Rotifers Is More Likely Due to Horizontal Gene Transfer Than to Meiotic Sex. Curr. Biol. 26, 723–732 (2016).

45. Mark Welch, D. B., Mark Welch, J. L. & Meselson, M. Evidence for degenerate tetraploidy in bdelloid rotifers. Proc. Natl. Acad. Sci. U. S. A. 105, 5145–5149 (2008).

46. Nowell, R. W. et al. Comparative genomics of bdelloid rotifers: Insights from desiccating and nondesiccating species. PLoS Biol. 16, e2004830 (2018).

47. Nowell, R. W. et al. Evolutionary dynamics of transposable elements in bdelloid rotifers. Elife 10, e63194 (2021).

48. Wilson, C. G., Nowell, R. W. & Barraclough, T. G. Cross-Contamination Explains “Inter and Intraspecific Horizontal Genetic Transfers” between Asexual Bdelloid Rotifers. Curr. Biol. 28, 2436–2444.e14 (2018).

49. Simion, P. et al. Homologous chromosomes in asexual rotifer *Adineta vaga* suggest automixis. bioRxiv 2020.06.16.155473 (2020) doi:10.1101/2020.06.16.155473.

50. Vakhrusheva, O. A. et al. Genomic signatures of recombination in a natural population of the bdelloid rotifer *Adineta vaga*. Nat. Commun. 11, 6421 (2020).

51. Gladyshev, E. A., Meselson, M. & Arkhipova, I. R. Massive horizontal gene transfer in bdelloid rotifers. Science 320, 1210–1213 (2008).

52. Signorovitch, A., Hur, J., Gladyshev, E. & Meselson, M. Allele sharing and evidence for sexuality in a mitochondrial clade of bdelloid rotifers. Genetics 200, 581–590 (2015).

53. Maynard Smith, J. Evolution: contemplating life without sex. Nature 324, 300–301 (1986).

54. Birky, C. W., Jr. Positively negative evidence for asexuality. J. Hered. 101 Suppl 1, S42–5 (2010).

55. Barron, G. L. Fungal parasites and predators of rotifers, nematodes, and other invertebrates. in Biodiversity of Fungi (eds. Mueller, G. M., Bills, G. F. & Foster, M. S.) 435–450 (Academic Press, 2004). doi:10.1016/B978-012509551-8/50022-2.

56. Wilson, C. G. & Sherman, P. W. Anciently asexual bdelloid rotifers escape lethal fungal parasites by drying up and blowing away. Science 327, 574–576 (2010).

57. Wilson, C. G. Desiccation-tolerance in bdelloid rotifers facilitates spatiotemporal escape from multiple species of parasitic fungi. Biol. J. Linn. Soc. Lond. 104, 564–574 (2011).

58. Wilson, C. G. & Sherman, P. W. Spatial and temporal escape from fungal parasitism in natural communities of anciently asexual bdelloid rotifers. Proc. Biol. Sci. 280, 20131255 (2013).

59. Engelstädter, J. Host-parasite coevolutionary dynamics with generalized success/failure infection genetics. Am. Nat. 185, E117–29 (2015).

60. Irazoqui, J. E., Urbach, J. M. & Ausubel, F. M. Evolution of host innate defence: insights from Caenorhabditis elegans and primitive invertebrates. Nat. Rev. Immunol. 10, 47–58 (2010).

61. Cogni, R., Cao, C., Day, J. P., Bridson, C. & Jiggins, F. M. The genetic architecture of resistance to virus infection in Drosophila. Mol. Ecol. 25, 5228–5241 (2016).

62. Bento, G. et al. The genetic basis of resistance and matching-allele interactions of a host-parasite system: The Daphnia magna-Pasteuria ramosa model. PLoS Genet. 13, e1006596 (2017).

63. Hudson, A. I., Fleming-Davies, A. E., Páez, D. J. & Dwyer, G. Genotype-by-genotype interactions between an insect and its pathogen. Journal of Evolutionary Biology 29, 2480–2490 (2016).

64. Ekroth, A. K. E., Gerth, M., Stevens, E. J., Ford, S. A. & King, K. C. Host genotype and genetic diversity shape the evolution of a novel bacterial infection. ISME J. 15, 2146–2157 (2021).

65. Fredericksen, M. et al. Infection phenotypes of a coevolving parasite are highly diverse, structured, and specific. Evolution (2021) doi:10.1111/evo.14323.

66. Boschetti, C., Pouchkina-Stantcheva, N., Hoffmann, P. & Tunnacliffe, A. Foreign genes and novel hydrophilic protein genes participate in the desiccation response of the bdelloid rotifer *Adineta ricciae*. J. Exp. Biol. 214, 59–68 (2011).

67. Yoshida, Y., Nowell, R. W., Arakawa, K. & Blaxter, M. Horizontal Gene Transfer in Metazoa: Examples and Methods. in Horizontal Gene Transfer: Breaking Borders Between Living Kingdoms (eds. Villa, T. G. & Viñas, M.) 203–226 (Springer International Publishing, 2019). doi:10.1007/978-3-030-21862-1_7.

68. Boschetti, C. et al. Biochemical diversification through foreign gene expression in bdelloid rotifers. PLoS Genet. 8, e1003035 (2012).

69. Richards, T. A. & Monier, A. A tale of two tardigrades. Proceedings of the National Academy of Sciences of the United States of America vol. 113 4892–4894 (2016).

70. Jaron, K. S. et al. Genomic Features of Parthenogenetic Animals. J. Hered. 112, 19–33 (2021).

71. Eyres, I. et al. Horizontal gene transfer in bdelloid rotifers is ancient, ongoing and more frequent in species from desiccating habitats. BMC Biol. 13, 90 (2015).

72. Rodriguez, F., Yushenova, I., DiCorpo, D. & Arkhipova, I. Bacterial N4-methylcytosine as an epigenetic mark in eukaryotic DNA. Research Square (2021) doi:10.21203/rs.3.rs-360382/v1.

73. Hecox-Lea, B. J. & Mark Welch, D. B. Evolutionary diversity and novelty of DNA repair genes in asexual Bdelloid rotifers. BMC Evol. Biol. 18, 177 (2018).

74. Harrison, E. & Brockhurst, M. A. Ecological and evolutionary benefits of temperate phage: What does or doesn’t kill you makes you stronger. Bioessays 39, (2017).

75. Hall, R. J., Whelan, F. J., McInerney, J. O., Ou, Y. & Domingo-Sananes, M. R. Horizontal Gene Transfer as a Source of Conflict and Cooperation in Prokaryotes. Front. Microbiol. 11, 1569 (2020).

76. Chou, S. et al. Transferred interbacterial antagonism genes augment eukaryotic innate immune function. Nature 518, 98–101 (2015).

77. Di Lelio, I. et al. Evolution of an insect immune barrier through horizontal gene transfer mediated by a parasitic wasp. PLOS Genetics 15, e1007998 (2019).

78. Verster, K. I. et al. Horizontal Transfer of Bacterial Cytolethal Distending Toxin B Genes to Insects. Molecular Biology and Evolution 36, 2105–2110 (2019).

79. Barron, G. L. A new genus, Rotiferophthora, to accommodate the Diheterospora-like endoparasites of rotifers. Can. J. Bot. 69, 494–502 (1991).

80. Segers, H. & Shiel, R. Tale of a sleeping beauty: a new and easily cultured model organism for experimental studies on bdelloid rotifers. Rotifera X 141–145 (2005).

81. Yoon, S. & Nam, D. Gene dispersion is the key determinant of the read count bias in differential expression analysis of RNA-seq data. BMC Genomics 18, 408 (2017).

82. Soneson, C. & Robinson, M. D. Bias, robustness and scalability in single-cell differential expression analysis. Nat. Methods 15, 255–261 (2018).

83. Jain, R., Rivera, M. C. & Lake, J. A. Horizontal gene transfer among genomes: the complexity hypothesis. Proc. Natl. Acad. Sci. U. S. A. 96, 3801–3806 (1999).

84. Gluck-Thaler, E. & Slot, J. C. Dimensions of Horizontal Gene Transfer in Eukaryotic Microbial Pathogens. PLoS Pathog. 11, e1005156 (2015).

85. Ricci, C. & Caprioli, M. Anhydrobiosis in bdelloid species, populations and individuals. Integr. Comp. Biol. 45, 759–763 (2005).

86. Mark Welch, D. B., Ricci, C. & Meselson, M. Bdelloid Rotifers: Progress in Understanding the Success of an Evolutionary Scandal. in Lost Sex 259–279 (Springer, Dordrecht, 2009). doi:10.1007/978-90-481-2770-2_13.

87. Hur, J. H., Van Doninck, K., Mandigo, M. L. & Meselson, M. Degenerate tetraploidy was established before bdelloid rotifer families diverged. Mol. Biol. Evol. 26, 375–383 (2009).

88. Sung, G.-H. et al. Phylogenetic classification of Cordyceps and the clavicipitaceous fungi. Stud. Mycol. 57, 5–59 (2007).

89. Herrero-Galán, E. et al. The insecticidal protein hirsutellin A from the mite fungal pathogen Hirsutella thompsonii is a ribotoxin. Proteins 72, 217–228 (2008).

90. Olombrada, M. et al. Fungal ribotoxins: Natural protein-based weapons against insects. Toxicon 83, 69–74 (2014).

91. Olombrada, M. et al. Characterization of a new toxin from the entomopathogenic fungus Metarhizium anisopliae: the ribotoxin anisoplin. Biological Chemistry 398, 135–142 (2017).

92. Maxwell Burroughs, A. & Aravind, L. RNA damage in biological conflicts and the diversity of responding RNA repair systems. Nucleic Acids Research 44, 8525–8555 (2016).

93. Brown, G. D. & Gordon, S. Immune recognition of fungal beta-glucans. Cell. Microbiol. 7, 471–479 (2005).

94. Hamilton, C. & Bulmer, M. S. Molecular antifungal defenses in subterranean termites: RNA interference reveals in vivo roles of termicins and GNBPs against a naturally encountered pathogen. Dev. Comp. Immunol. 36, 372–377 (2012).

95. Liu, T. et al. Structural and biochemical insights into an insect gut-specific chitinase with antifungal activity. Insect Biochem. Mol. Biol. 119, 103326 (2020).

96. Boller, T. Antimicrobial Functions of the Plant Hydrolases, Chitinase and ß-1,3-Glucanase. in Mechanisms of Plant Defense Responses (eds. Fritig, B. & Legrand, M.) 391–400 (Springer Netherlands, 1993). doi:10.1007/978-94-011-1737-1_124.

97. Aktuganov, G. et al. Wide-range antifungal antagonism of Paenibacillus ehimensis IB-X-b and its dependence on chitinase and beta-1,3-glucanase production. Canadian Journal of Microbiology 54, 577–587 (2008).

98. McTaggart, S. J., Conlon, C., Colbourne, J. K., Blaxter, M. L. & Little, T. J. The components of the Daphnia pulex immune system as revealed by complete genome sequencing. BMC Genomics 10, 175 (2009).

99. Viljakainen, L. Evolutionary genetics of insect innate immunity. Briefings in Functional Genomics 14, 407–412 (2015).

100. Leulier, F. & Lemaitre, B. Toll-like receptors--taking an evolutionary approach. Nature Reviews Genetics 9, 165–178 (2008).

101. Pukkila-Worley, R., Ausubel, F. M. & Mylonakis, E. Candida albicans infection of Caenorhabditis elegans induces antifungal immune defenses. PLoS Pathog. 7, e1002074 (2011).

102. Spoel, S. H., Johnson, J. S. & Dong, X. Regulation of tradeoffs between plant defenses against pathogens with different lifestyles. Proc. Natl. Acad. Sci. U. S. A. 104, 18842–18847 (2007).

103. Palmer, W. J. & Jiggins, F. M. Comparative Genomics Reveals the Origins and Diversity of Arthropod Immune Systems. Molecular Biology and Evolution 32, 2111–2129 (2015).

104. Peters, A. D. & Lively, C. M. The Red Queen and Fluctuating Epistasis: A Population Genetic Analysis of Antagonistic Coevolution. Am. Nat. 154, 393–405 (1999).

105. Sasaki, A., Hamilton, W. D. & Ubeda, F. Clone mixtures and a pacemaker: new facets of Red-Queen theory and ecology. Proc. Biol. Sci. 269, 761–772 (2002).

106. Gandon, S. & Otto, S. P. The evolution of sex and recombination in response to abiotic or coevolutionary fluctuations in epistasis. Genetics 175, 1835–1853 (2007).

107. Salathé, M., Kouyos, R. D., Regoes, R. R. & Bonhoeffer, S. Rapid parasite adaptation drives selection for high recombination rates. Evolution 62, 295–300 (2008).

108. Salathé, M., Kouyos, R. D. & Bonhoeffer, S. The state of affairs in the kingdom of the Red Queen. Trends Ecol. Evol. 23, 439–445 (2008).

109. Davis, H. A new callidina: With the result of experiments on the desiccation of rotifers. Mon. Microsc. J. 9, 201–209 (1873).

110. Örstan, A. The trouble with Adineta vaga (Davis, 1873): a common rotifer that cannot be identified (Rotifera: Bdelloidea: Adinetidae). Zootaxa 4830, zootaxa.4830.3.8 (2020).

111. Ricci, C. Culturing of some bdelloid rotifers. Hydrobiologia 112, 45–51 (1984).

112. Mark Welch, D. B. & Meselson, M. Oocyte nuclear DNA content and GC proportion in rotifers of the anciently asexual Class Bdelloidea. Biol. J. Linn. Soc. Lond. 79, 85–91 (2003).

113. Pouchkina-Stantcheva, N. N. et al. Functional divergence of former alleles in an ancient asexual invertebrate. Science 318, 268–271 (2007).

114. Barron, G. L. Fungal parasites of bdelloid rotifers: Diheterospora. Can. J. Bot. 63, 211–222 (1985).

115. Jenkins, D. G. & Underwood, M. O. Zooplankton may not disperse readily in wind, rain, or waterfowl. in Rotifera VIII: A Comparative Approach 15–21 (Springer Netherlands, 1998). doi:10.1007/978-94-011-4782-8_3.

116. Fontaneto, D., Barraclough, T. G., Chen, K., Ricci, C. & Herniou, E. A. Molecular evidence for broad-scale distributions in bdelloid rotifers: everything is not everywhere but most things are very widespread. Mol. Ecol. 17, 3136–3146 (2008).

117. Fontaneto, D. et al. Cryptic diversity in the genus Adineta Hudson & Gosse, 1886 (Rotifera: Bdelloidea: Adinetidae): a DNA taxonomy approach. Hydrobiologia 662, 27–33 (2011).

118. McCallum, H. et al. Breaking beta: deconstructing the parasite transmission function. Philos. Trans. R. Soc. Lond. B Biol. Sci. 372, (2017).

119. Pujol, N. et al. Anti-fungal innate immunity in C. elegans is enhanced by evolutionary diversification of antimicrobial peptides. PLoS Pathog. 4, e1000105 (2008).

120. Zhang, W. et al. Comparative transcriptomic analysis of immune responses of the migratory locust, Locusta migratoria, to challenge by the fungal insect pathogen, Metarhizium acridum. BMC Genomics 16, 867 (2015).

121. Lu, H.-L. & St Leger, R. J. Insect Immunity to Entomopathogenic Fungi. Adv. Genet. 94, 251–285 (2016).

122. Grabherr, M. G. et al. Full-length transcriptome assembly from RNA-Seq data without a reference genome. Nat. Biotechnol. 29, 644–652 (2011).

123. Haas, B. J. et al. De novo transcript sequence reconstruction from RNA-seq using the Trinity platform for reference generation and analysis. Nat. Protoc. 8, 1494–1512 (2013).

124. Quast, C. et al. The SILVA ribosomal RNA gene database project: improved data processing and web-based tools. Nucleic Acids Res. 41, D590–6 (2013).

125. Dobin, A. et al. STAR: ultrafast universal RNA-seq aligner. Bioinformatics 29, 15–21 (2013).

126. Andrews, S. FastQC: A quality-control tool for high-throughput sequence data. (2015).

127. Ewels, P., Magnusson, M., Lundin, S. & Käller, M. MultiQC: summarize analysis results for multiple tools and samples in a single report. Bioinformatics 32, 3047–3048 (2016).

128. Patro, R., Duggal, G., Love, M. I., Irizarry, R. A. & Kingsford, C. Salmon provides fast and bias-aware quantification of transcript expression. Nat. Methods 14, 417–419 (2017).

129. Love, M. I., Huber, W. & Anders, S. Moderated estimation of fold change and dispersion for RNA-seq data with DESeq2. Genome Biol. 15, 550 (2014).

130. Benjamini, Y. & Hochberg, Y. Controlling the False Discovery Rate: A Practical and Powerful Approach to Multiple Testing. J. R. Stat. Soc. Series B Stat. Methodol. 57, 289–300 (1995).

131. Li, H. Minimap2: pairwise alignment for nucleotide sequences. Bioinformatics 34, 3094–3100 (2018).

132. Simão, F. A., Waterhouse, R. M., Ioannidis, P., Kriventseva, E. V. & Zdobnov, E. M. BUSCO: assessing genome assembly and annotation completeness with single-copy orthologs. Bioinformatics 31, 3210–3212 (2015).

133. UniProt Consortium. UniProt: a worldwide hub of protein knowledge. Nucleic Acids Res. 47, D506–D515 (2019).

134. El-Gebali, S. et al. The Pfam protein families database in 2019. Nucleic Acids Res. 47, D427–D432 (2019).

135. Altschul, S. F., Gish, W., Miller, W., Myers, E. W. & Lipman, D. J. Basic local alignment search tool. J. Mol. Biol. 215, 403–410 (1990).

136. Petersen, T. N., Brunak, S., von Heijne, G. & Nielsen, H. SignalP 4.0: discriminating signal peptides from transmembrane regions. Nat. Methods 8, 785–786 (2011).

137. Krogh, A., Larsson, B., von Heijne, G. & Sonnhammer, E. L. Predicting transmembrane protein topology with a hidden Markov model: application to complete genomes. J. Mol. Biol. 305, 567–580 (2001).

138. Jones, P. et al. InterProScan 5: genome-scale protein function classification. Bioinformatics 30, 1236–1240 (2014).

139. Bryant, D. M. et al. A Tissue-Mapped Axolotl De Novo Transcriptome Enables Identification of Limb Regeneration Factors. Cell Rep. 18, 762–776 (2017).

140. Buchfink, B., Xie, C. & Huson, D. H. Fast and sensitive protein alignment using DIAMOND. Nat. Methods 12, 59–60 (2015).

141. Koutsovoulos, G. et al. No evidence for extensive horizontal gene transfer in the genome of the tardigrade Hypsibius dujardini. Proceedings of the National Academy of Sciences of the United States of America vol. 113 5053–5058 (2016).

142. Nguyen, L.-T., Schmidt, H. A., von Haeseler, A. & Minh, B. Q. IQ-TREE: a fast and effective stochastic algorithm for estimating maximum-likelihood phylogenies. Mol. Biol. Evol. 32, 268–274 (2015).

143. Kalyaanamoorthy, S., Minh, B. Q., Wong, T. K. F., von Haeseler, A. & Jermiin, L. S. ModelFinder: fast model selection for accurate phylogenetic estimates. Nat. Methods 14, 587–589 (2017).

144. Minh, B. Q. et al. IQ-TREE 2: New models and efficient methods for phylogenetic inference in the genomic era. Mol. Biol. Evol. (2020) doi:10.1093/molbev/msaa015.

145. Emms, D. M. & Kelly, S. OrthoFinder: phylogenetic orthology inference for comparative genomics. Genome Biol. 20, 238 (2019).

146. Han, J. et al. The genome of the marine monogonont rotifer *Brachionus plicatilis*: Genome-wide expression profiles of 28 cytochrome P450 genes in response to chlorpyrifos and 2-ethyl-phenanthrene. Aquat. Toxicol. 214, 105230 (2019).

